# Meta-analyses reveal support for the Social Intelligence Hypothesis

**DOI:** 10.1101/2024.05.15.594271

**Authors:** Elizabeth M. Speechley, Benjamin J. Ashton, Yong Zhi Foo, Leigh W. Simmons, Amanda R. Ridley

## Abstract

The Social Intelligence Hypothesis (SIH) is one of the leading explanations for the evolution of cognition. Since its inception a vast body of literature investigating the predictions of the SIH has accumulated, using a variety of methodologies and species. However, the generalisability of the hypothesis remains unclear. To gain an understanding of the robustness of the SIH as an explanation for the evolution of cognition, we systematically searched the literature for studies investigating the predictions of the SIH. Accordingly, we compiled 103 studies with 584 effect sizes from 17 taxonomic orders. We present the results of four meta-analyses which reveal support for the SIH across interspecific, intraspecific and developmental studies. However, effect sizes did not differ significantly between the cognitive or sociality metrics used, taxonomy or testing conditions. Thus, support for the SIH is similar across studies using neuroanatomy and cognitive performance, those using broad categories of sociality, group size and social interactions, across taxonomic groups, and for tests conducted in captivity or the wild. Overall, our meta-analyses support the SIH as an evolutionary and developmental explanation for cognitive variation.

## I. INTRODUCTION

### (1) The Social Intelligence Hypothesis

Despite over a century of investigation and widespread interest, the evolution of cognition remains the focus of substantial debate (van Horik & Emery, 2011). Cognition can be defined as the mental processes by which animals collect, retain and use information from their environment to guide their behaviour (Shettleworth, 2001). The Social Intelligence Hypothesis (SIH) has emerged as one of the leading theories to explain the evolution of cognition, predicting that the demands of social interactions are an important selective factor (Dunbar, 1998; Jolly, 1966; Humphrey, 1976; Chance & Mead, 1953). The link between cognition and sociality dates back to Darwin (1871), however the SIH was initially proposed by Chance & Mead (1953) and Jolly (1966), and then formalised by Humphrey (1976). The SIH has since taken various forms, such as the Machiavellian Intelligence Hypothesis (Byrne & Whiten, 1988) and the Social Brain Hypothesis (Dunbar, 1998), but the central tenet remains the same: that cognition has evolved in response to the demands of living in complex social groups, in which individuals interact in different contexts with multiple different individuals, and often repeatedly with the same individuals over time (Freeberg, Dunbar & Ord, 2012). Maintaining and coordinating multiple relationships, monitoring both intra- and intergroup individuals, and recognising suitable cooperative partners, are a few of the many cognitive demands required of social animals that are hypothesised to require a high degree of informational processing (Dunbar, 1998; Jolly, 1966; Humphrey, 1976; Chance & Mead, 1953).

Most research has investigated the relationship between cognition and sociality from an evolutionary perspective by comparing neuroanatomy and sociality across species (e.g. Dunbar, 1992; DeCasien & Higham, 2019; Shultz & Dunbar, 2006). However, an increasing number of studies are taking an intraspecific approach to the study of cognition (Ashton, Thornton & Ridley, 2018*b*; Thornton & Lukas, 2012), exploring how the social environment may influence cognitive performance among individuals of the same species. Additionally, some studies have manipulated rearing conditions experimentally to explore how the social environment may impact cognitive development (Bannier, Tebbich & Taborsky, 2017; Carducci & Jakob, 2000; Riley *et al*., 2018; Fischer *et al*., 2015). However, evidence for the SIH based on these approaches is varied and conflicting. For example, in some comparative studies sociality predicts variation in relative brain size (Shultz & Dunbar, 2007), whereas in others it does not (Kverková *et al*., 2018; DeCasien, Williams & Higham, 2017). Likewise, some intraspecific studies show conflicting results using the same cognitive trait in the same species: for example, in brown rats (*Rattus norvegicus*) one study detected a positive relationship between social housing and spatial memory (Heimer-McGinn *et al*., 2020), whereas another detected no relationship (Harris, D’Eath & Healy, 2009). Consequently, the relationship between the social environment and cognition, at both the evolutionary and developmental level, remains unclear.

Given the widespread interest in the SIH, a quantitative analysis of existing literature testing the hypothesis is timely and much needed. Meta-analyses allow us to synthesise existing tests and quantitatively analyse support for a hypothesis (Nakagawa *et al*., 2017). Therefore, we attempt to fill this knowledge gap by presenting four separate phylogenetically controlled meta-analyses (*sensu* Nakagawa & Santos, 2012) of the SIH in non-human animals based on four study categories (Fig. 1). Specifically, we test (*a*) evidence for the SIH as an evolutionary hypothesis by analysing comparative interspecific tests split into two subcategories depending on how these interspecific tests were conducted (i.e. using species averages or groups of individuals from different species). We also investigate (*b*) whether there is evidence for support of the SIH in intraspecific studies, and (*c*) the mechanistic underpinnings of the hypothesis by investigating developmental studies testing the effect of the early-life social environment on cognitive development. Additionally, we conduct moderator analyses to examine factors which may be affecting heterogeneity in the sociality– cognition relationship across studies (see Section I.2 and Fig. 1). Finally, to analyse the robustness of our results we estimate publication bias according to year of publication, and biases towards studies with small sample sizes and large effect sizes.

**Fig. 1.**
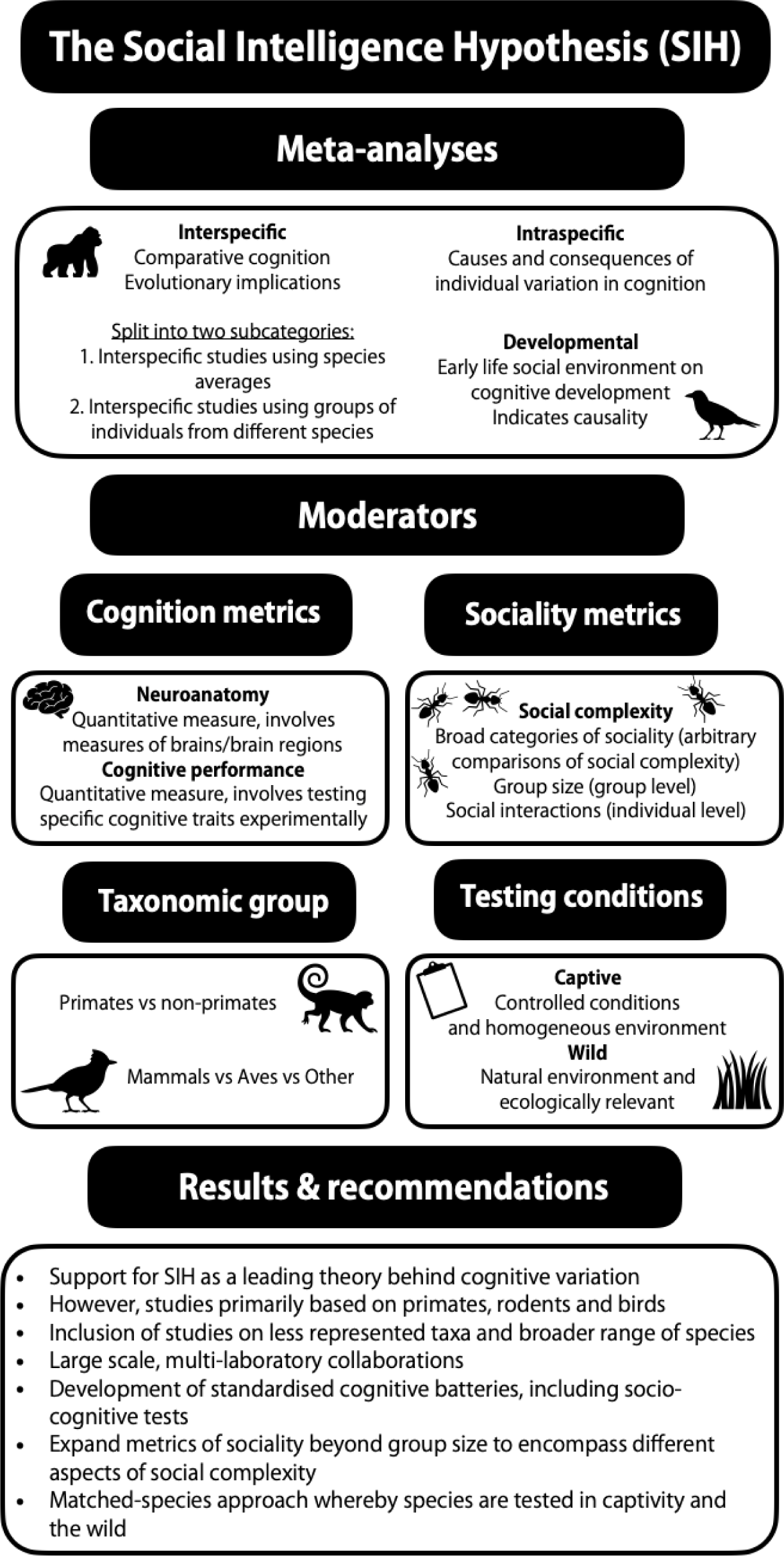
Schematic summarising the four meta-analyses, the moderators, our findings and recommendations for future research.

### (2) Disentangling support for the Social Intelligence Hypothesis

Meta-analysis also gives us a tool to identify the moderators that account for heterogeneity in effect sizes across studies by testing whether effect sizes vary across moderator levels through meta-regression models (Nakagawa *et al*., 2023). Here, we examine the following moderators: cognition metrics (neuroanatomy or cognitive performance), sociality metric, taxonomic group and testing conditions (wild or captive) (Fig. 1), each of which is discussed in detail below.

Regardless of whether studies are tested intra- or interspecifically, an impressive array of neuroanatomical proxies and behavioural metrics have been used to test the predictions of the SIH. Studies of the SIH began by using brain morphology [originally based on variants of Jerison’s encephalisation quotient (Jerison, 1973) as a measure of relative brain size] as a proxy for cognition, including measures of relative brain size (Shultz & Dunbar, 2006, 2007) and/or brain regions such as the neocortex (Dunbar, 1992, 1995), amygdala (Noonan *et al*., 2014; Sallet *et al*., 2011), cerebrum (Sakai *et al*., 2011), cerebellum (Sallet *et al*., 2011; Sakai *et al*., 2011), telencephalon (Burish, Kueh & Wang, 2004), the high vocal centre (Gil *et al*., 2006), and white and grey brain matter (Meguerditchian *et al*., 2021). However, investigations of the relationship between sociality and multiple measures of neuroanatomy have generated conflicting results (Todorov *et al*., 2019; Deaner, Nunn & van Schaik, 2000; Joffe & Dunbar, 1997; DeCasien & Higham, 2019; Arsznov & Sakai, 2012; Seid & Junge, 2016; Swanson *et al*., 2012), indicating that the SIH may only apply to certain brain regions. For example, a study on the cleaner fish (*Labroides dimindiatus*) found a positive relationship between the density of cleaner fish and the size of the diencephalon and telencephalon, whereas the size of the mesencephalon, rhombencephalon and brain stem were unrelated to cleaner fish density (Triki *et al*., 2019). Cognitive performance has also been measured using a variety of procedures to test specific cognitive traits, including associative and reversal learning (Riley *et al*., 2018; Liedtke & Schneider, 2017; Ashton *et al*., 2018*a*; Bond, Kamil & Balda, 2007; Bannier *et al*., 2017; Meagher *et al*., 2015), innovation (Ashton, Thornton & Ridley, 2019; Morand-Ferron & Quinn, 2011; Dean *et al*., 2011; Liker & Bókony, 2009), transitive inference (Bond, Kamil & Balda, 2003; Bond, Wei & Kamil, 2010; Maclean, Merritt & Brannon, 2008), spatial learning/memory (Langley *et al*., 2018a,*b*; Riley *et al*., 2017; Ashton *et al*., 2018*a*; Costanzo, Bennett & Lutermann, 2009; Machatschke *et al*., 2011), quantity discrimination (Kelly, 2016), and inhibitory control (Maclean *et al*., 2013, 2014; Johnson-Ulrich & Holekamp, 2020; Reddy *et al*., 2015; Amici, Aureli & Call, 2008). However, tests also report conflicting results within the same cognitive trait (Wang *et al*., 2018; Williams *et al*., 2001). For example, studies manipulating housing group size of house mice (*Mus musculus)* found different patterns of association between group size and cognition depending on the cognitive test used (Wang *et al*., 2018; Williams *et al*., 2001; Kogan, Frankland & Silva, 2000; Smith *et al*., 2018). There is also debate as to whether neuroanatomical measures provide robust proxies of cognition (Healy & Rowe, 2007; Logan *et al*., 2018; Healy & Rowe, 2013), especially given that certain taxonomic groups exhibit similar cognitive skills despite vastly different brain structure (Güntürkün & Bugnyar, 2016). Therefore, we attempt to answer this question by exploring whether support for the SIH differs for studies using direct behavioural tests of cognition compared to those using neuroanatomical proxies, by including cognitive metric as a moderator in our analysis. Due to the debate regarding the validity of neuroanatomical proxies, we predict direct cognitive measures to have larger effect sizes compared to indirect measures such as neuroanatomy.

In contrast to the diversity of cognitive measures used to explore the SIH, relatively less attention has been given to the social component of the hypothesis. Sociality is broadly defined as the tendency to live as part of a group with clear organisation of social interactions (American Psychological Association, 2022). SIH studies typically quantify sociality by categorising species as solitary or non-solitary, or based on their mating system (Arsznov & Sakai, 2013; De Meester, Huyghe & Van Damme, 2019), the temporal variation in social grouping (e.g. temporary aggregations or permanent groups) (Bond *et al*., 2010), or by social subunits or social hierarchies (Krasheninnikova *et al*., 2013). However, by far the most commonly used sociality metric is social group size (Dunbar, 1992, 1995; Fichtel, Dinter & Kappeler, 2020; Lindenfors, 2005; Lindenfors, Nunn & Barton, 2007; Navarrete *et al*., 2016; Louail *et al*., 2019; Sobrero *et al*., 2016; Keverne, Martel & Nevison, 1996). Group size is a valuable social metric because it gives the largest potential number of conspecifics an individual can interact with and can be readily applied to both inter- and intra-specific tests. However, group size assumes that all individuals within the same group will experience the same social pressures, when realistically, individuals in social groups may not be interacting equally (Morrison *et al*., 2020). The presence of more individuals also does not necessarily correspond to greater social complexity, as individuals may not differentiate between conspecifics (Bergman & Beehner, 2015). With the rise of intraspecific tests, increasing interest has been devoted to individual-based measures of sociality based on social network analysis. These studies also reported equivocal support for the relationship between sociality and cognition, depending on the metric tested (Aplin *et al*., 2012; Wascher, 2015; Kulahci *et al*., 2016; Kulahci, Ghazanfar & Rubenstein, 2018; Preiszner *et al*., 2015; Ash, Ziegler & Colman, 2020; Berhane & Gazes, 2020; Cowl & Shultz, 2017; Graham, 2011; Kudo & Dunbar, 2001; Lehmann & Dunbar, 2009; Lewis & Barton, 2006; Pasquaretta *et al*., 2014). Overall, it remains unclear whether support for the SIH holds across group- and individual-based sociality metrics. Therefore, we test for differences in effect sizes across sociality metrics to investigate whether effect sizes vary based on studies using broad measures of sociality (e.g. social system), group-based measures of sociality (e.g. group size) or individual-based measures of sociality (e.g. social network metrics).

The SIH began to gain traction in the 1990s after a positive correlation between neocortex size and group size was identified in anthropoid primates (Dunbar, 1992). The relationship between group size and brain size/neocortex size in anthropoid primates has since been replicated in several other studies (Dunbar, 1995; Barton, 1996; Walker *et al*., 2006; Dunbar & Shultz, 2007), which inevitably led to the expansion of the SIH beyond primates. Since its inception, support for the SIH has been found in various species of mammals (Wang *et al*., 2018; Johnson-Ulrich & Holekamp, 2020; Fox, Muthukrishna & Shultz, 2017; Sakai *et al*., 2011; Dunbar & Bever, 1998; Borrego & Gaines, 2016), birds (Beauchamp & Fernández-Juricic, 2004; Kulahci *et al*., 2016; Langley *et al*., 2018*b*; Aplin *et al*., 2012; Boogert, Farine & Spencer, 2014; Morand-Ferron & Quinn, 2011; Lipkind *et al*., 2002; Ashton *et al*., 2018*a*; Speechley *et al*., 2024), fish (Fischer *et al*., 2015; Triki *et al*., 2019; Brandão, Braithwaite & Goncalves-de-Freitas, 2015; Stanbrook *et al*., 2020; Leris & Reader, 2016; Ausas *et al*., 2019), and invertebrates (Liedtke & Schneider, 2017; Kamhi *et al*., 2016; Ott & Rogers, 2010; Amador-Vargas *et al*., 2015; Seid & Junge, 2016). However, support for the SIH has not been consistent among or within taxa (Templeton, Kamil & Balda, 1999; Iwaniuk & Arnold, 2004; Kverková *et al*., 2018; Forss *et al*., 2016; DeCasien *et al*., 2017; Fedorova, Evans & Byrne, 2017), with many papers reporting conflicting results, even within the same species, depending on the metric of sociality or cognition used (Fox *et al*., 2017; Kamhi *et al*., 2016; Sakai *et al*., 2016; Reader, Hager & Laland, 2011; DeCasien & Higham, 2019).

Consequently, we examine whether the hypothesis applies beyond primates by testing for differences in effect sizes between primates and non-primates. If the effects are applicable beyond primates, we should find no differences in effect sizes across taxa. If the hypothesis is applicable only to primates, we should find a larger effect size for primates and a smaller, non-significant effect for non-primates. We predict studies on non-primate taxa to have similar effect sizes compared to those on primate taxa.

Tests of the relationship between sociality and cognition have been conducted under a range of conditions, including both in the wild and in captivity, but little is known about the potential impact of testing conditions on the data generated. Studies of the SIH, particularly neuroanatomical studies, are often carried out on captive individuals (e.g. Noonan *et al*., 2014; Sallet *et al*., 2011; Sandel, MacLean & Hare, 2011; Petit *et al*., 2015), but studies on wild and wild-caught captive individuals are becoming increasingly common (Kulahci *et al*., 2018; Langley *et al*., 2018*b*; Ashton *et al*., 2018*a*; Speechley *et al*., 2024), and both methodologies have received criticism. For example, testing in the wild introduces many confounding factors, such as previous experience and motivational inconsistencies, that may influence results if not controlled for experimentally or statistically [see Rowe & Healy (2014), Morand-Ferron, Cole & Quinn (2016) and Boogert *et al*. (2018) for innovative solutions]. Additionally, captive individuals are often exposed to stress due to isolation during experimental testing – particularly for social species (Cazakoff, Johnson & Howland, 2010; Hobson *et al*., 2019) – which is likely to have consequences for brain function and cognition (Hobson *et al*., 2019; Weiss *et al*., 2004). Given this debate, we also investigate whether effect sizes vary across wild and captive testing conditions.

## II. METHODS

### (1) Literature search

We undertook a systematic literature search in *Scopus* and *Web of Science* (WOS). Key words were combined with Boolean operators (e.g. AND, OR) and wildcards (*) to account for alternate word endings, quotation marks were used where we were interested in a particular combination of words (e.g. “social complexity”) and parentheses were used for nested searches. The search was split into two parts, the first part of the search focused on key words related to sociality, whilst the second part of the search focused on cognition and measures of neuroanatomy. The final search string, formatted for *Scopus* was: ALL (sociality OR “social complexity” OR “social group*” OR “group size” OR “social network*” OR “social structure” OR machiavellian OR “social intelligence” OR “social brain hypothesis”) AND (cognition OR “brain size*” OR “brain region*”OR neocortex OR hippocampus OR “inhibitory control”) AND INDEXTERMS (animal OR animalia OR bird OR primate OR mammal OR insect* OR fish OR reptile) AND NOT TITLE (“human”). ‘Inhibitory control’ was specifically included in the search to avoid exclusion of some relevant SIH papers.

This search restricted results to non-human animal papers by using the index terms and excluding papers with ‘human’ in the title and key words. Furthermore, the search was limited to articles, conference papers, books and book chapters written in English. This generated 5567 results on 22/06/2021. The WOS search generated 2163 results on 22/06/2021, giving us a combined total of 7730 publications. Once duplicates were removed (353), this left a total of 7377 papers (Fig. 2). We searched for grey literature (e.g. unpublished post-graduate dissertations) using an identical search string from passed theses in EBSCOhost. Our search results are summarised according to PRISMA guidelines (Liberati *et al*., 2009) in Fig. 2.

**Fig. 2.**
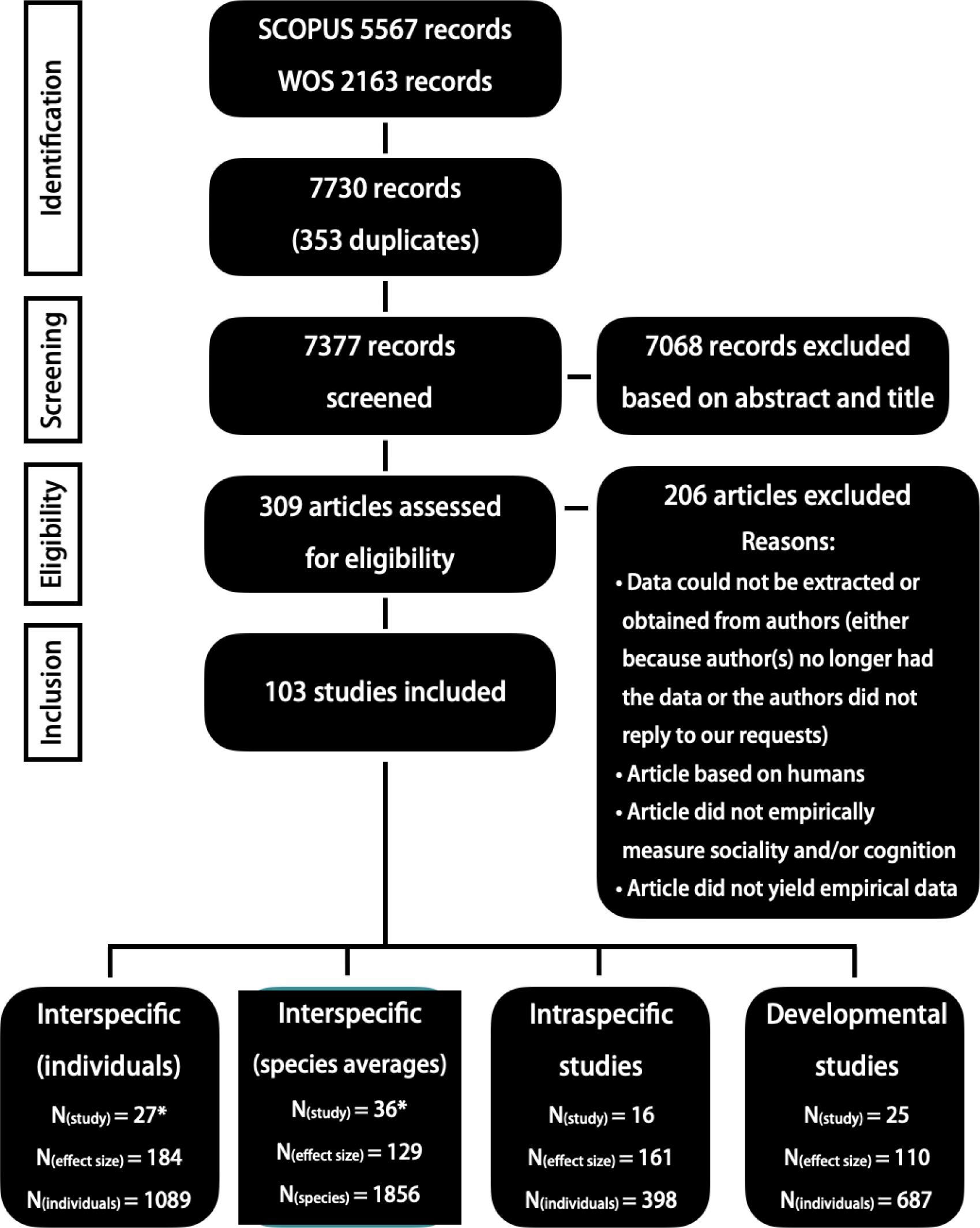
PRISMA diagram showing systematic search process including data filtering (with filtering justification). The final number of studies, effect sizes and individuals in each of the four meta-analyses is given. Interspecific (individual) corresponds to comparative interspecific tests based on groups of individuals from different species, whereas interspecific (species averages) corresponds to comparative interspecific tests based on species averages. Intraspecific studies involved within-species tests, and developmental studies include studies testing the effect of early life environment on cognitive development. *One study was included in two meta-analyses: interspecific (individual) and interspecific (species averages).

### (2) Inclusion criteria

We used three criteria to determine if a paper was suitable for inclusion: the paper needed to (1) focus on non-human species and (2) have an empirical measure of sociality and cognition. We rejected studies that did not yield empirical data such as reviews or theoretical studies. In addition, the paper (3) must also contain extractable data (effect size and sample size values or statistics that allowed us to infer values). For papers that did not contain extractable data we contacted authors for their original data sets. These papers were excluded if we could not contact the authors or if the data were unavailable. We initially screened the data for these inclusion criteria by reading the title and abstract of each paper, which narrowed the search down from 7377 to 309 articles. The remaining 309 articles were further filtered to 103 studies during the data extraction process based on screening of the full text and availability of data sets (see online Supporting Information, Table S1 for list of all papers included in meta-analyses). It is important to note that our search captured some studies that did not directly set out to test the SIH, despite having a social measure which was compared to cognitive performance or neuroanatomy. However, we included them as their results were relevant to our hypothesis. The systematic assessment of data was conducted by E.M.S., but 10% of papers were assessed by a second participant (P.R., see Acknowledgements) to assess potential observer bias.

### (3) Effect size extraction/calculation

We used Pearson’s *r* as our measure of effect size. We extracted *r*, means and variance estimates (SD, SE), and proportional data from published studies. For studies without reported means and variance, we extracted *F* values and *t* statistics and converted to *r* values using the formulae in Lipsey & Wilson (2001). For studies that reported multiple measures of cognition across time, we took the mean and variance between the largest time interval (shortest to longest time). Likewise, for multifactorial studies that contained a control group and groups at different social densities, we focused on the difference between the lowest and highest density groups. Where means and dispersion data were provided only in figures, these values were extracted using the R package *metaDigitise* (Pick, Nakagawa & Noble, 2019). As a normal distribution is required for analysis, Pearson’s *r* was converted to Fischer’s *Z* (*Zr*) which is not bounded (Hedges & Olkin, 1985). Results were then back-transformed to Pearson’s *r* to facilitate interpretation (i.e. positive *r* equates to a positive relationship between sociality and cognition).

### (4) Moderator variables and coding of papers

For every effect size, we recorded the publication title and year of publication. Each study was categorised into one of four groups: (*a*) interspecific (individuals), when the study compared two or more species by testing groups of individuals from different species, (*b*) interspecific (species averages), when the study compared two or more species using species averages; (*c*) intraspecific, when the study tested individuals within a species; and (*d*) developmental, when individuals were tested during development, prior to reaching sexual maturity. Each of these experimental categories was tested *via* a separate meta-analysis (Fig. 1). We recorded the following moderator variables: (*a*) taxonomic group at the class, order and species level (where possible); (*b*) testing conditions (wild, captive, unknown); (*c*) sociality metrics (e.g. group size); and (*d*) cognition metrics. Cognition and sociality metrics were then broadly categorised to facilitate comparisons. Specifically, for cognition these categories included studies based on neuroanatomical proxies and those based on cognitive tests. Sociality was categorised based on how the study measured sociality, namely whether they used broad metrics of sociality (e.g. social system: solitary species *versus* aggregations, temporal variation in social grouping or the existence of social subunits/social hierarchies), group size, or individual social interactions (e.g. frequency of affiliation, agonism, vocalisations; Fig. 1).

### (5) Phylogenies

We used R v.4.2.1 (R Core Team, 2022) to create a phylogeny for each meta-analysis using the *rotl* package (Michonneau, Brown & Winter, 2016) (Fig. 3). *rotl* uses the Interactive Tree of Life database (ITOL; Hinchliff *et al*., 2015) to generate a tree using data from the National Centre for Biotechnology Information Taxonomy database (Wheeler *et al*., 2006). Once a tree was created for each meta-analysis, we generated a variance–covariance matrix amongst species using the topology of the tree (i.e. the evolutionary relationships among the species without branch lengths) and incorporated this matrix as a random effect. Phylogeny not only controls for non-independence in data coming from the same species, but also provides an indication of whether the effect sizes vary across species.

**Fig. 3.**
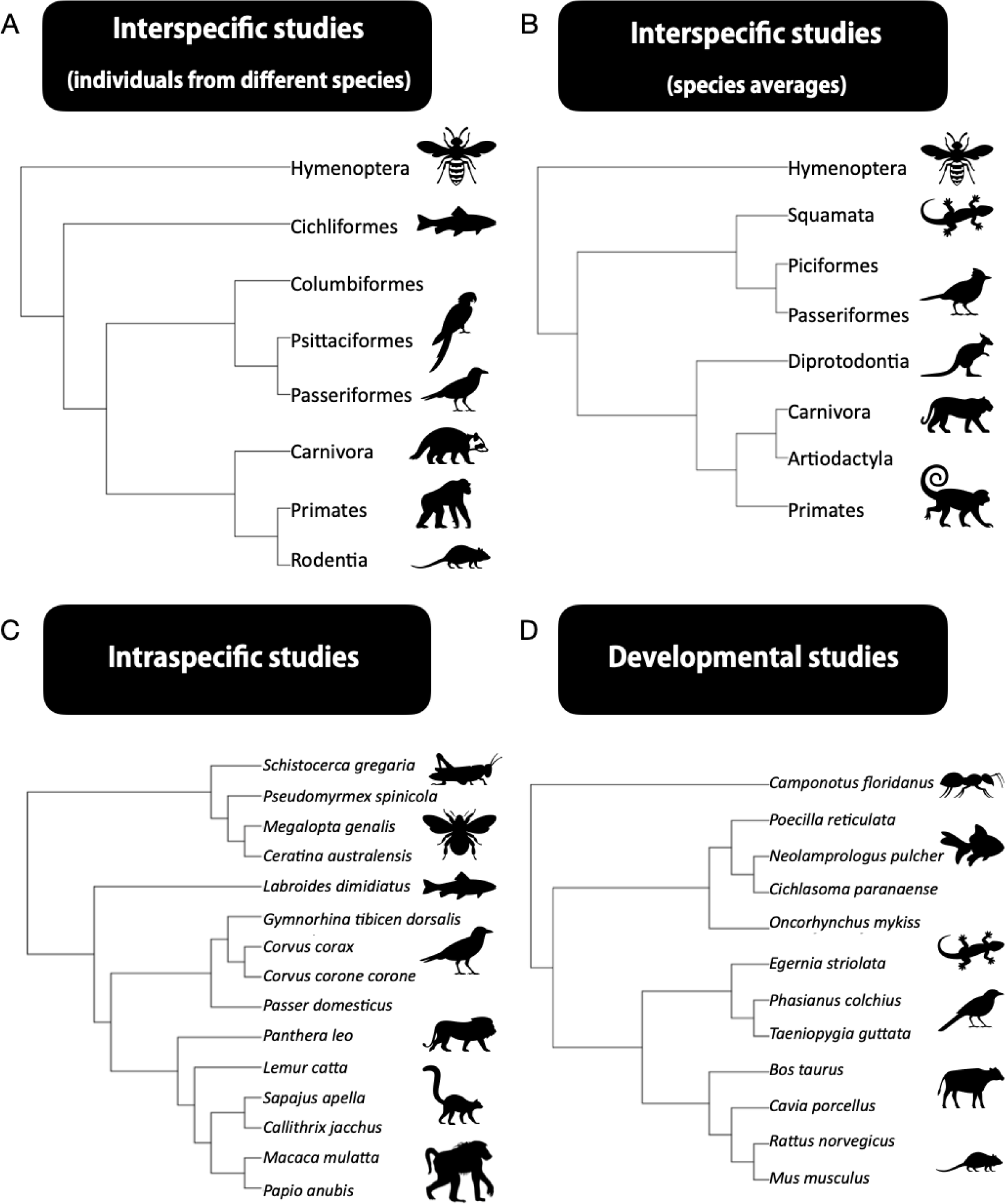
Phylogenetic trees for taxa included in each meta-analysis: (A) interspecific studies based on individual tests (*N* = 8 orders); (B) interspecific studies based on species averages (*N* = 8 orders); (C) intraspecific studies (*N* = 15 species); and (D) developmental studies (*N* = 12 species). A and B show taxonomic orders and C and D show species included in the meta-analyses.

### (6) Meta-analysis

We ran multilevel meta-analyses using linear mixed models via the *metafor* package (Nakagawa & Poulin, 2012; Viechtbauer, 2010) in R v.4.2.1 (R Core Team, 2022). Linear mixed models allow us to control for non-independence in the data due to multiple effect sizes extracted from the same study, effect size identity which allows the results to vary across effect sizes, multiple effect sizes from the same species, or shared ancestry amongst species. Therefore, study identity, effect size identity, species identity, and phylogeny were included as potential random effects in each model. We first ran the models without any random effects to generate a heterogeneity (*I^2^*) value using the *rma* function in *metafor* (Viechtbauer, 2010). The *I^2^* value indicates the percentage of variation that is due to heterogeneity rather than chance (Higgins & Thompson, 2002; Higgins *et al*., 2003). A high *I^2^* (>75%) means there is large variation between effect sizes and thus moderator analyses are necessary.

For each meta-analysis we began by checking the relative importance of each random effect based on the *I^2^* of the random variables using an intercept-only model using restricted maximum likelihood estimation (REML) with the four random variables listed above. We determined the relative importance of the random effect of each variable in comparison to each other by computing the heterogeneity statistic (*I^2^*) based on the above model using the *rma* function in *metafor* (Viechtbauer, 2010). Each random factor that accounted for a substantial proportion of the total *I^2^* (i.e. *I^2^*>0%) was included in the meta-analysis. Overall effect sizes were tested for each analysis by running an intercept-only model using REML estimation with the selected random variable/s. All models were conducted with robust variance estimation to account for non-independence in errors. Finally, *Zr* results were back-transformed to *r* for interpretation purposes.

### (7) Moderator selection

Before conducting moderator analyses, we evaluated multicollinearity using Goodman and Kruskal’s *t* measure of association between categorical predictor variables using the package *GoodmanKruskal* (Upton, 2005). Variables with correlation values above 60% were removed from analyses to achieve a balance between avoiding the inclusion of highly correlated predictor values and excluding too many predictors. Additionally, we calculated generalised variance inflation factor (GVIF) values (allowing us to assess multicollinearity between categorical predictors) using the *vif* function from the *metafor* package (Viechtbauer, 2010) and converted this to variance inflation factor (VIF) values to check for multicollinearity. Predictors with VIF values of <2 were included in the final models (VIF >2 for pairs of continuous predictors was considered a level of correlation too high to include the predictors together in the analysis) (Fox & Monette, 1992).

We also assessed the distribution of effect sizes within each level of a moderator, to ensure the difference between the minimum and maximum effect sizes in a moderator variable was approximately 1:10 (Harrell Jr., Lee & Mark, 1996). Where this 1:10 ratio was not met, levels within the moderator were condensed or the moderator was removed from analysis. Specifically, taxonomy was condensed into primates *versus* other taxonomic groups in interspecific studies using groups of individuals from different species and interspecific studies using species averages due to lack of effect sizes for non-primate taxa (Table 1). However, we were able to expand taxonomy to mammals, birds and other groups for the meta-analysis of developmental studies (no primate taxa were tested in developmental studies).

**Table 1.**
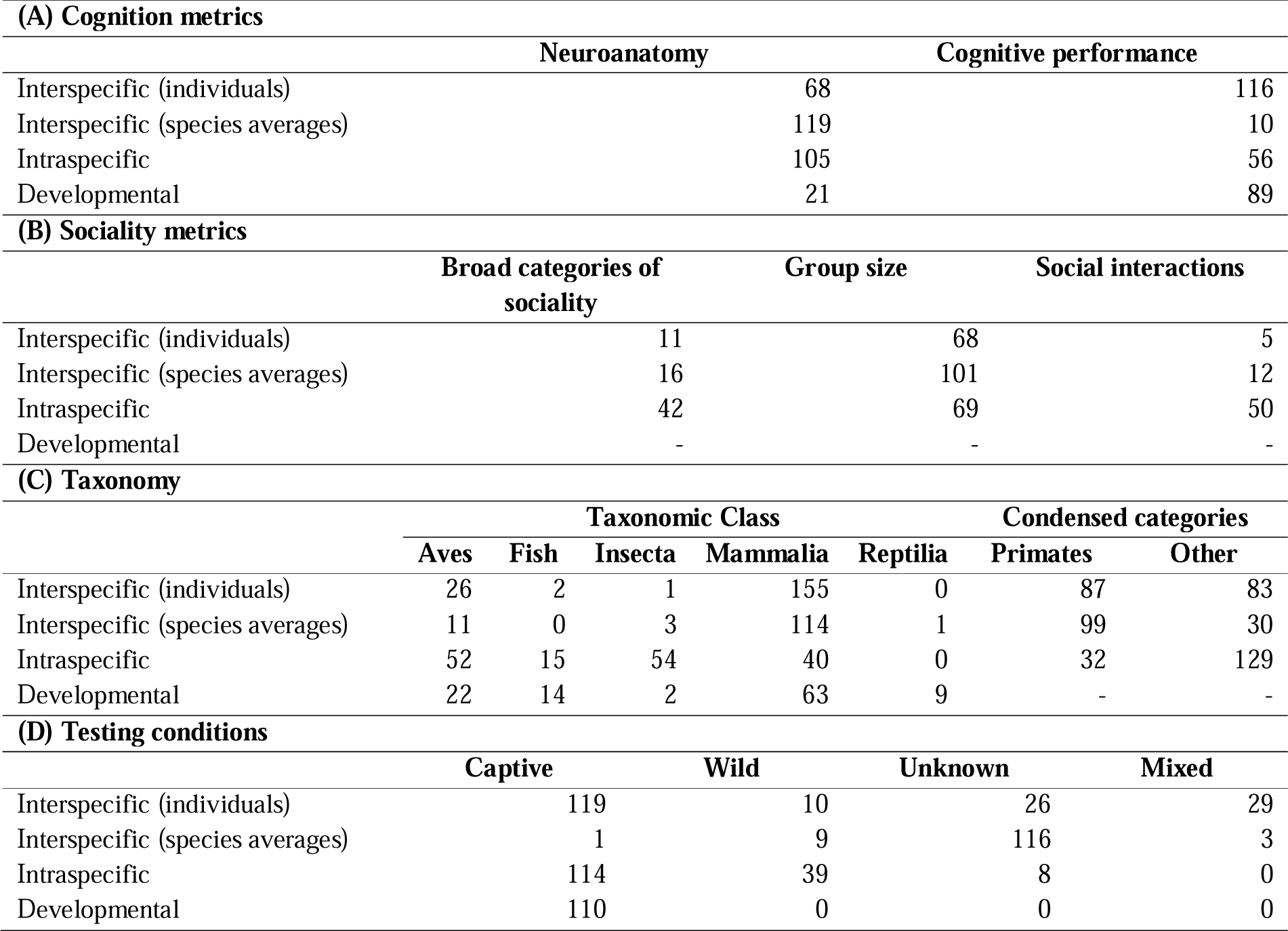
Distribution of effect sizes for each moderator: (A) cognition metrics, (B) sociality metrics, (C) taxonomy and (D) testing conditions across each of the four meta-analyses.

Cognition and sociality metrics and taxonomic group were included as moderators in all analyses (except for developmental studies where sociality metrics were excluded because all studies used group size to measure sociality) (Table 1). Taxonomy was excluded from intraspecific studies as it was highly correlated with other moderators (sociality and testing condition, see Table S2). Testing condition was excluded from interspecific studies using groups of individuals from different species because it was highly correlated with cognitive performance (see Table S3). It was also removed from interspecific studies using species averages as most effect sizes came from unknown origins (Table 1) and because the overwhelming majority (96%) of studies were conducted in captivity for developmental studies.

### (8) Moderator analysis

To conduct moderator analyses, meta-regression models were run using the *rma.mv* function in *metafor* (Nakagawa & Poulin, 2012; Viechtbauer, 2010). We first ran single-factor models without the intercept using the REML estimation, including only one categorical moderator as a fixed factor and the designated random factors. This allowed us to obtain parameter estimates of each level in each factor after controlling for the random factors. Additionally, we calculated a marginal *R^2^* value for each of these moderators to determine the proportion of variance the moderator explained (Table S4). We then conducted automated model selection (Grueber *et al*., 2011) using the package *MuMIn* (Bartoń, 2022) to identify moderators that remained in the final model. This was based on Akaike’s Information Criterion with sample size correction (AICc) obtained from the maximum likelihood (ML) estimation. Model selection was run using only the effect sizes that had no missing data to ensure that the AICc values of the different models were comparable (Nakagawa & Freckleton, 2010). We created all possible candidate models from the relevant random factors and moderator variables. The top model set was defined as all models within two AICc units of the best model (i.e. the model with the lowest AICc value). If the null model was retained within a top model set, we considered none of our variables to be good predictors of data patterns. Additionally, if a moderator was not included in the top model set, this suggested that there was no significant difference in the effect sizes between our moderator variables.

### (9) Publication bias

We tested for a publication bias using multilevel meta-regression which can model heterogeneity and non-independence (Nakagawa *et al*., 2022). Specifically, we examined evidence for a time-lag bias, which is the tendency for some studies to be published faster than others depending on the direction and magnitude of the results, with larger effects published first (Jennions & Møller, 2002). We also test for whether studies with small sample size and large effects are more likely to be published *via* an effective sample size calculation (Nakagawa *et al*., 2022). To obtain an overall estimate adjusted for publication bias, we first ran a single-factor model, with year of publication (centred on 0) and effective sample size 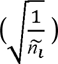 as fixed factors (where 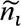 is the *i*th sample size) and the same random factors as described above (Nakagawa *et al*., 2022). If the intercept was non-significant, we used 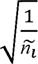 to check for significance of funnel asymmetry (small-sample-size effects). Alternatively, if the intercept was significant, we used 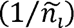 to obtain an overall estimate adjusted for publication bias.

## III. RESULTS

### (1) Overall effects

We found a significant positive relationship between sociality and cognition in three out of four meta-analyses after accounting for their respective random effects (see Table S5 for *I^2^* values). In the following text, *N* is given as number of effect sizes with number of articles in parentheses. For interspecific studies based on individual tests, the relationship between sociality and cognition was not statistically significant (*r* = 0.16, CI = –0.03/0.35, *P* = 0.09, *N* = 184(27), Table 2A, Fig. 4A). However, we found a significant relationship between sociality and cognition in interspecific studies using species averages (*r* = 0.15, CI = 0.01/0.30, *P* = 0.04, *N* = 129(36), Table 2B, Fig. 5A), intraspecific studies (*r* = 0.23, CI = 0.04/0.42, *P* = 0.02, *N* = 161(16), Table 2C, Fig. 6A) and developmental studies (*r* = 0.24, CI = 0.11/0.38, *P* = 0.00, *N* = 110(25); Table 2D, Fig. 7A).

**Fig. 4.**
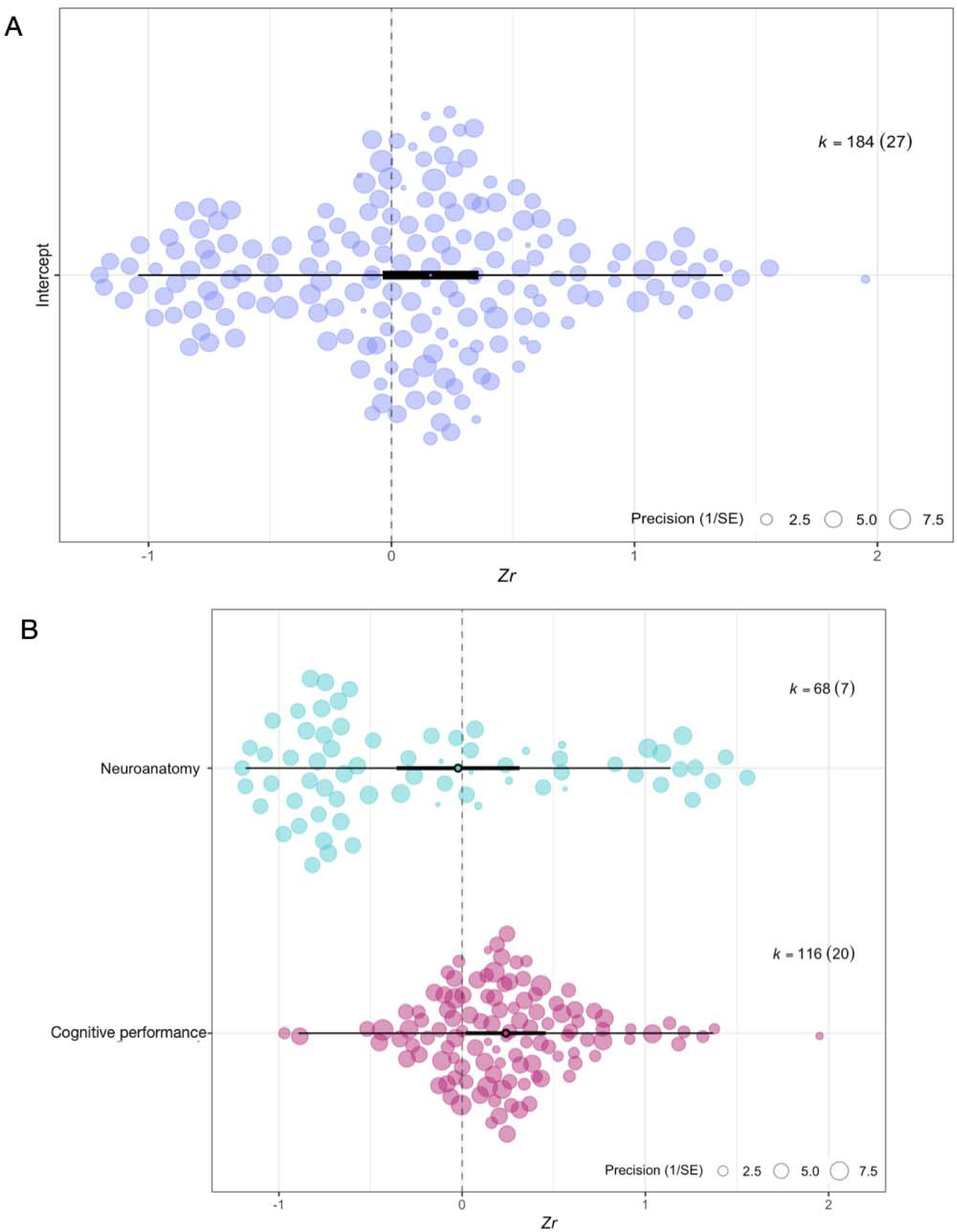
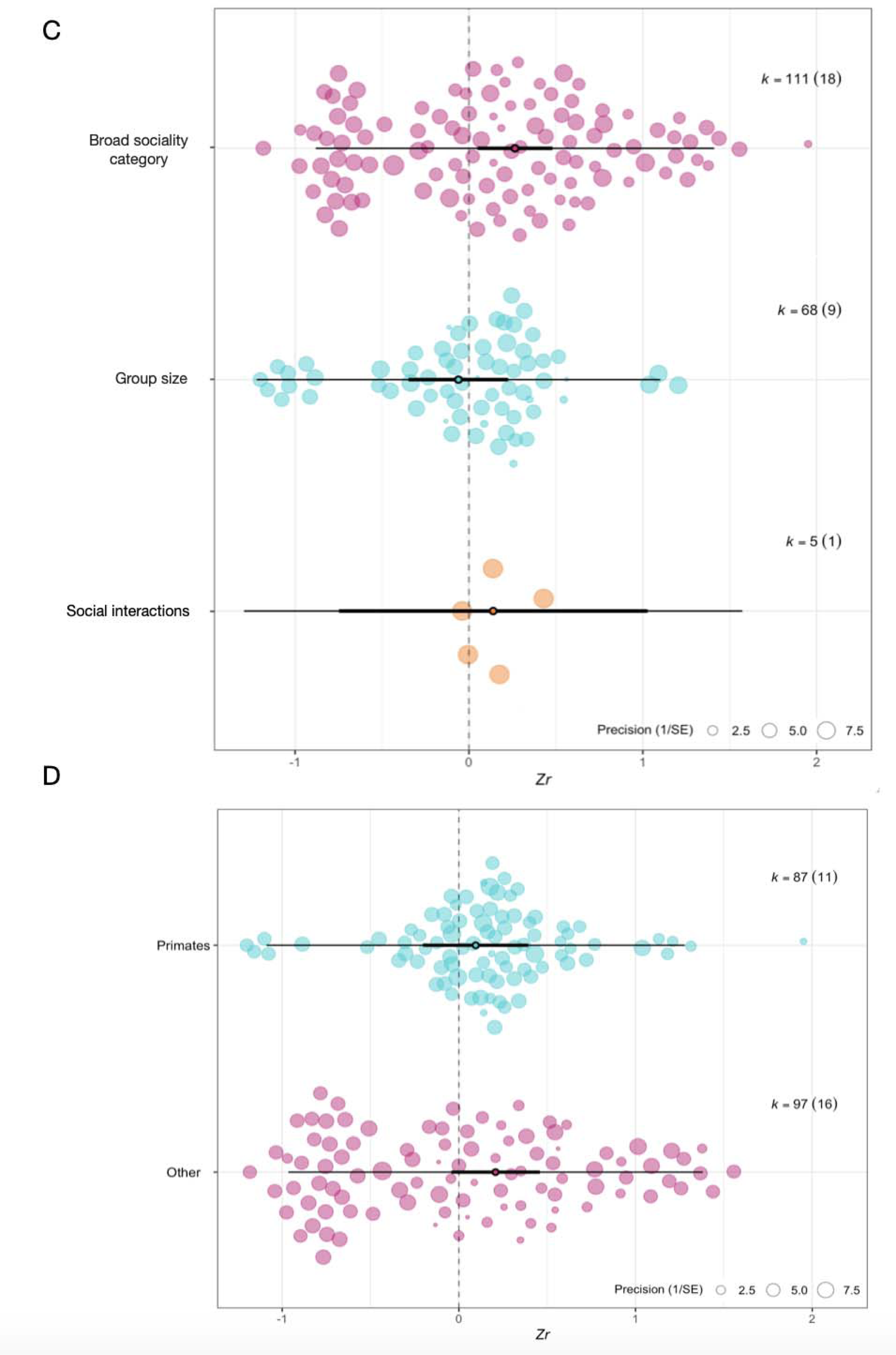
Orchard plots of (A) overall effect size (*Zr*) estimates centred on zero for interspecific studies based on individual tests. Orchard plots are also shown for moderators in these studies, including (B) cognitive metric used, (C) sociality metric, and (D) taxonomic group. The thick central box represents 95% confidence intervals, the central point is the mean and the thin line represents point estimates. The size of the points represents the precision (1/SE), whereby larger points represent larger sample sizes and thus greater precision.

**Fig. 5.**
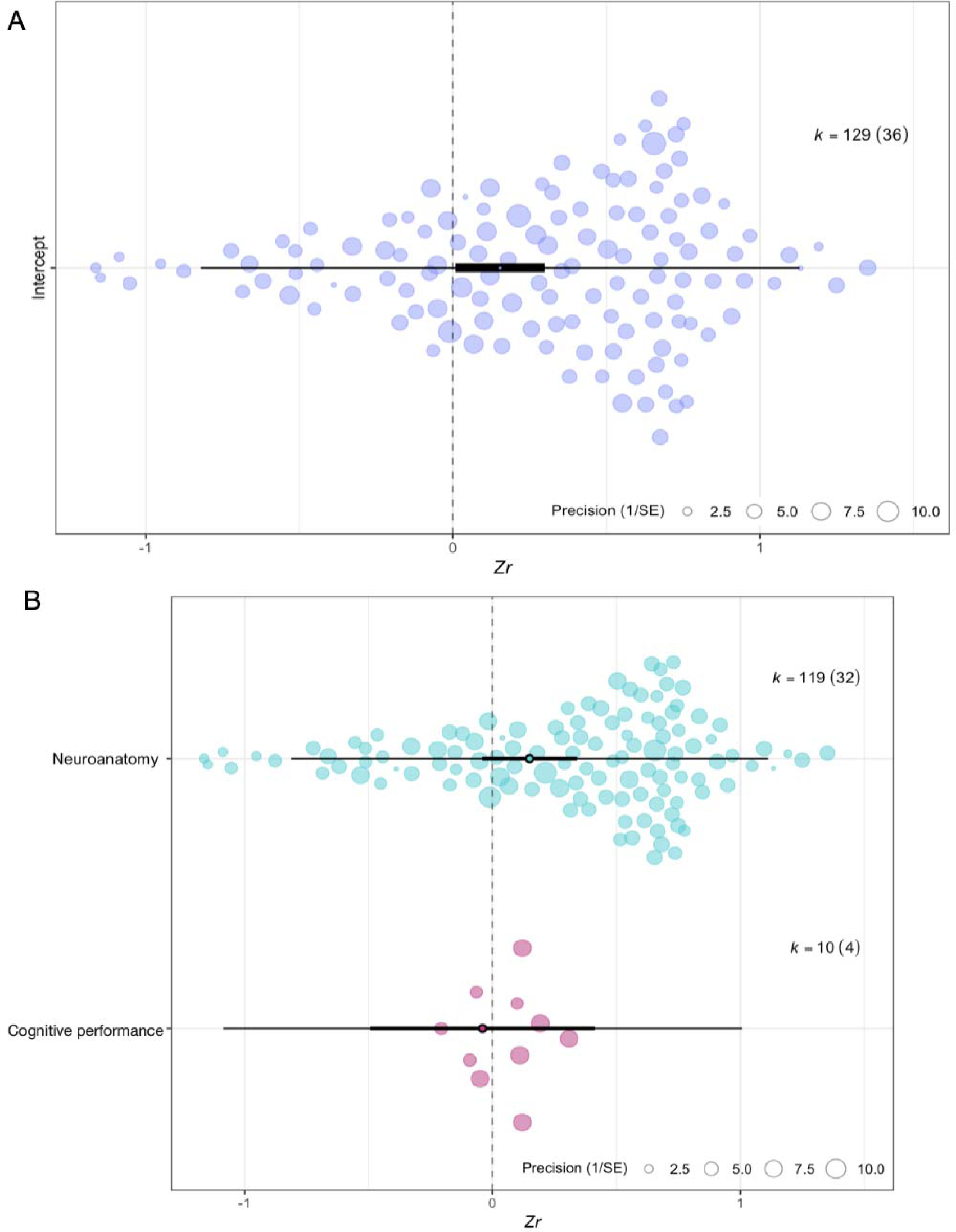
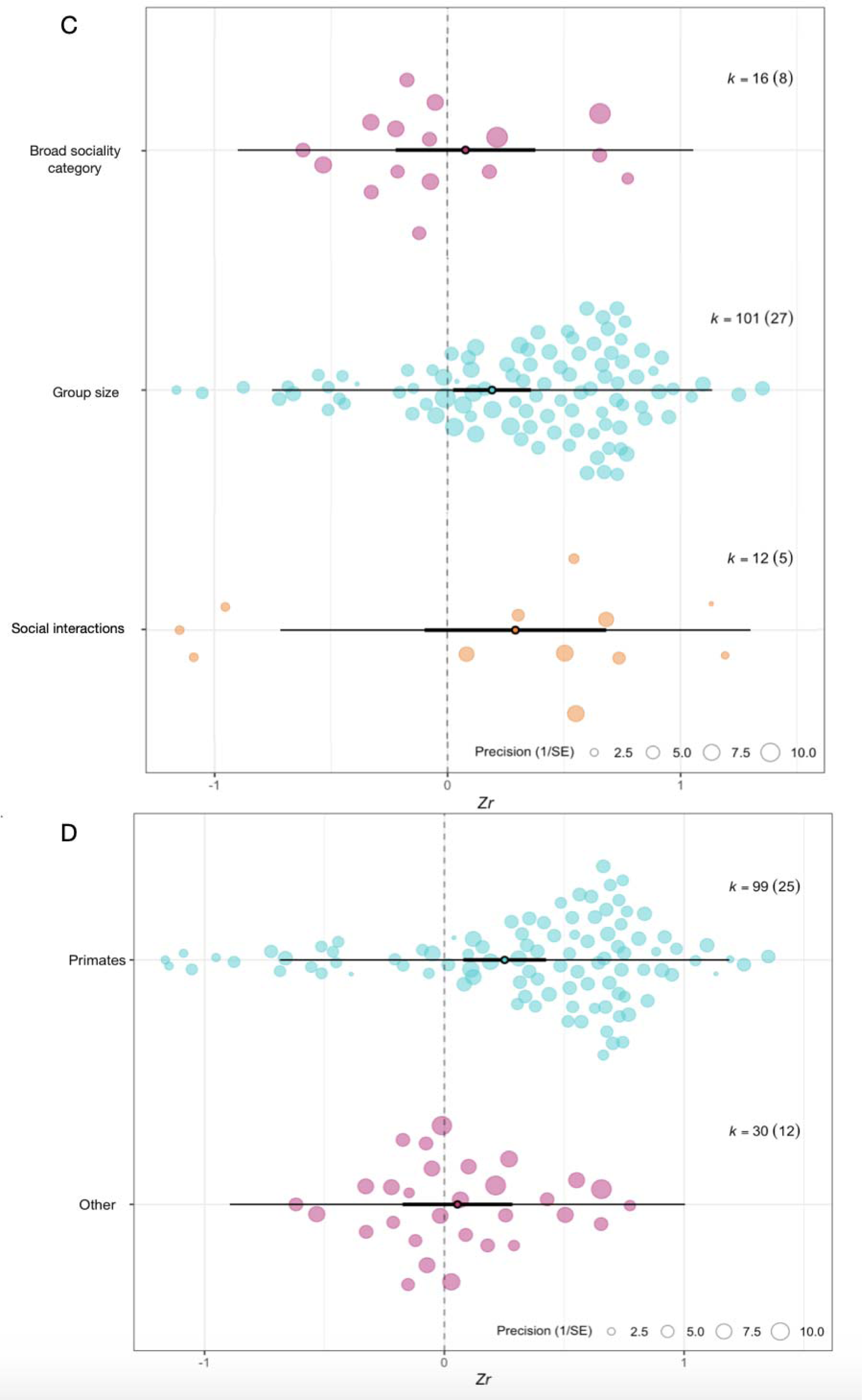
Orchard plots of (A) overall effect size (*Zr*) estimates centred on zero for interspecific studies based on species averages (confidence intervals are positive and did not intersect 0). Orchard plots are also shown for moderators in these studies, including (B) cognition metrics used, (C) sociality metrics, and (D) taxonomic group. The thick central box represents 95% confidence intervals, the central point is the mean and the thin line represents point estimates. The size of the points represents the precision (1/SE), whereby larger points represent larger sample sizes and thus greater precision.

**Fig. 6.**
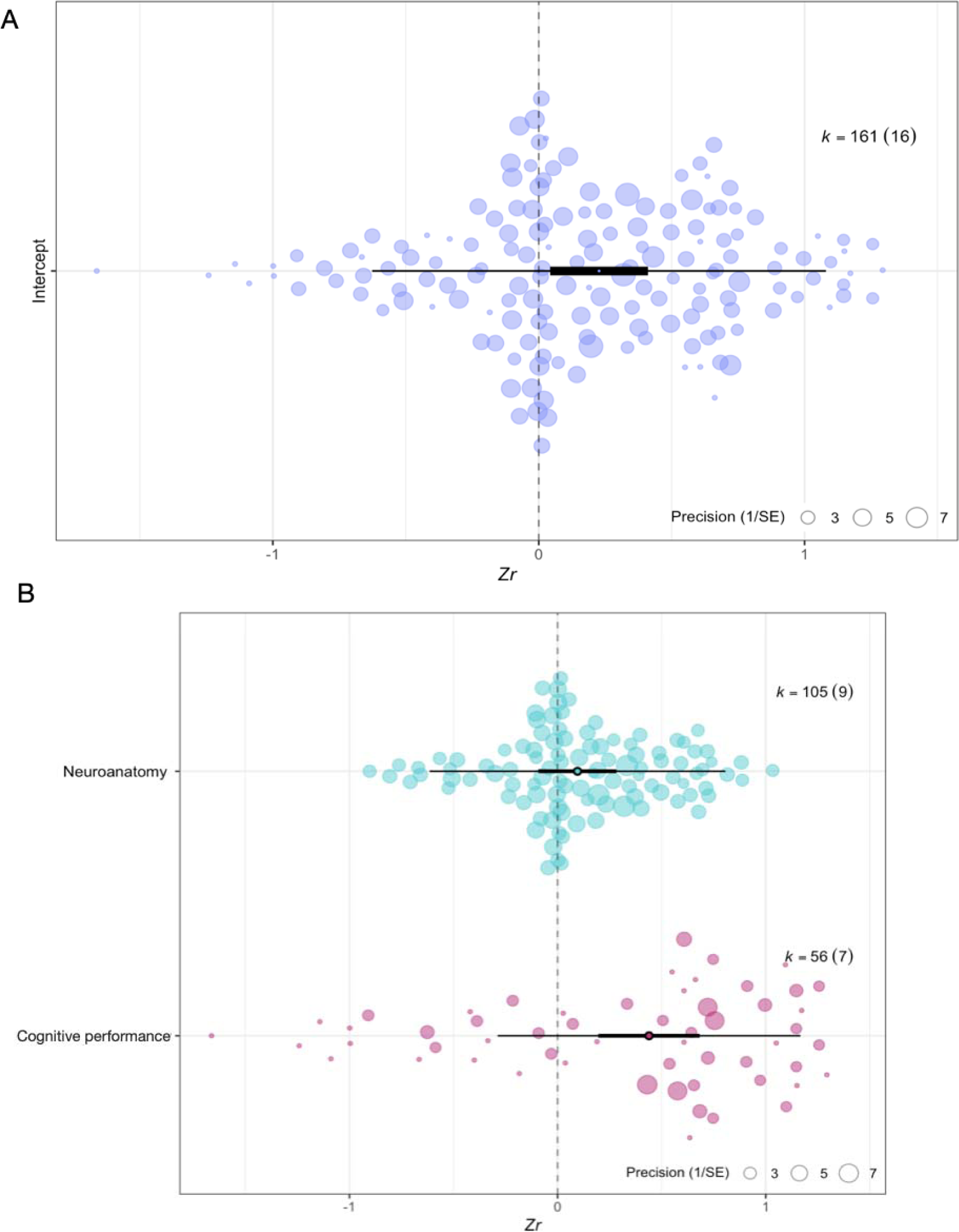
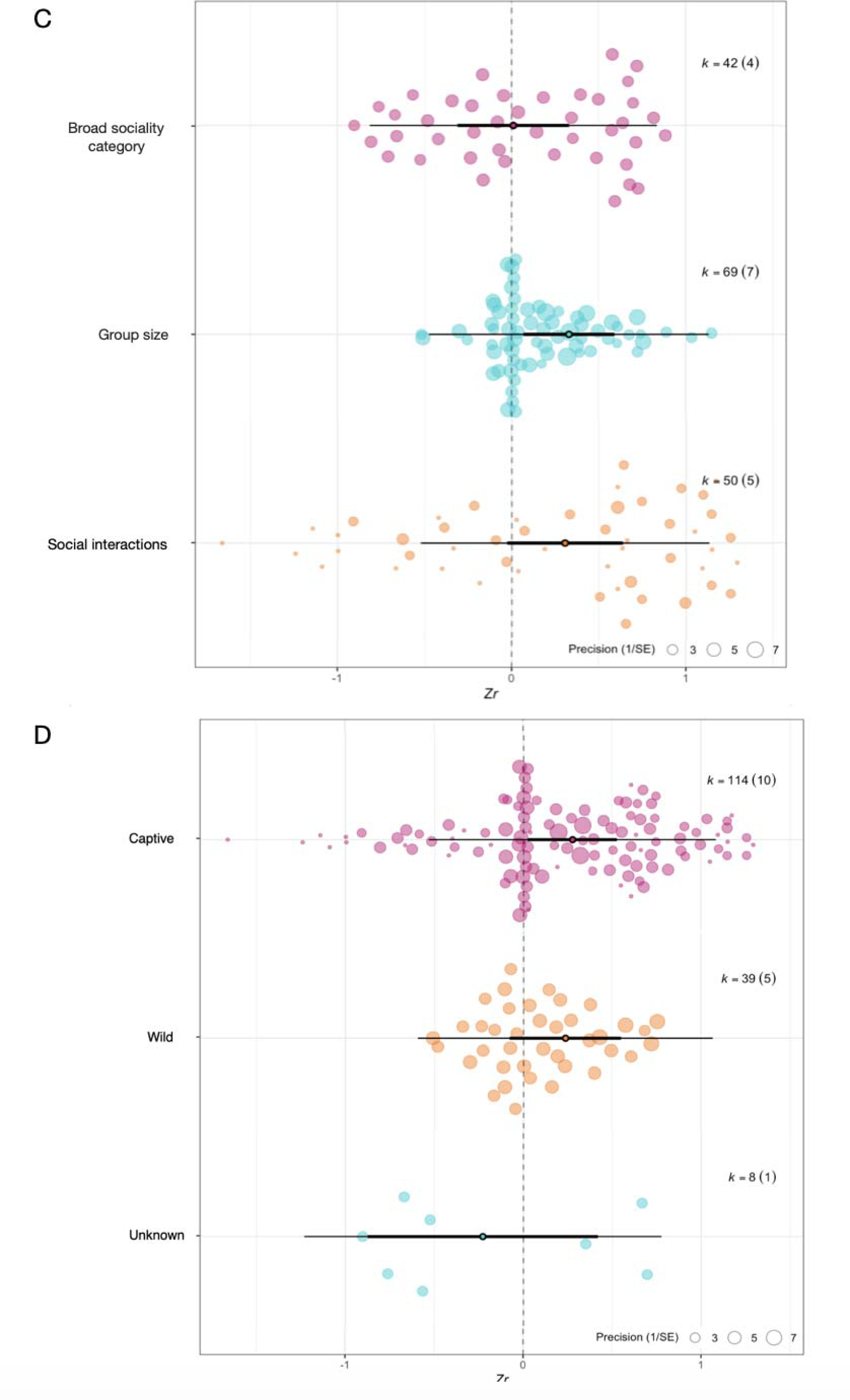
Orchard plots of (A) overall effect size (*Zr*) estimates centred on zero for intraspecific studies (confidence intervals are positive and did not intersect 0). Orchard plots are also shown for moderators in these studies, including (B) cognition metrics used, (C) sociality metrics, and (D) testing conditions. The thick central box represents 95% confidence intervals, the central point is the mean and the thin line represents point estimates. The size of the points represents the precision (1/SE), whereby larger points represent larger sample sizes and thus greater precision.

**Fig. 7.**
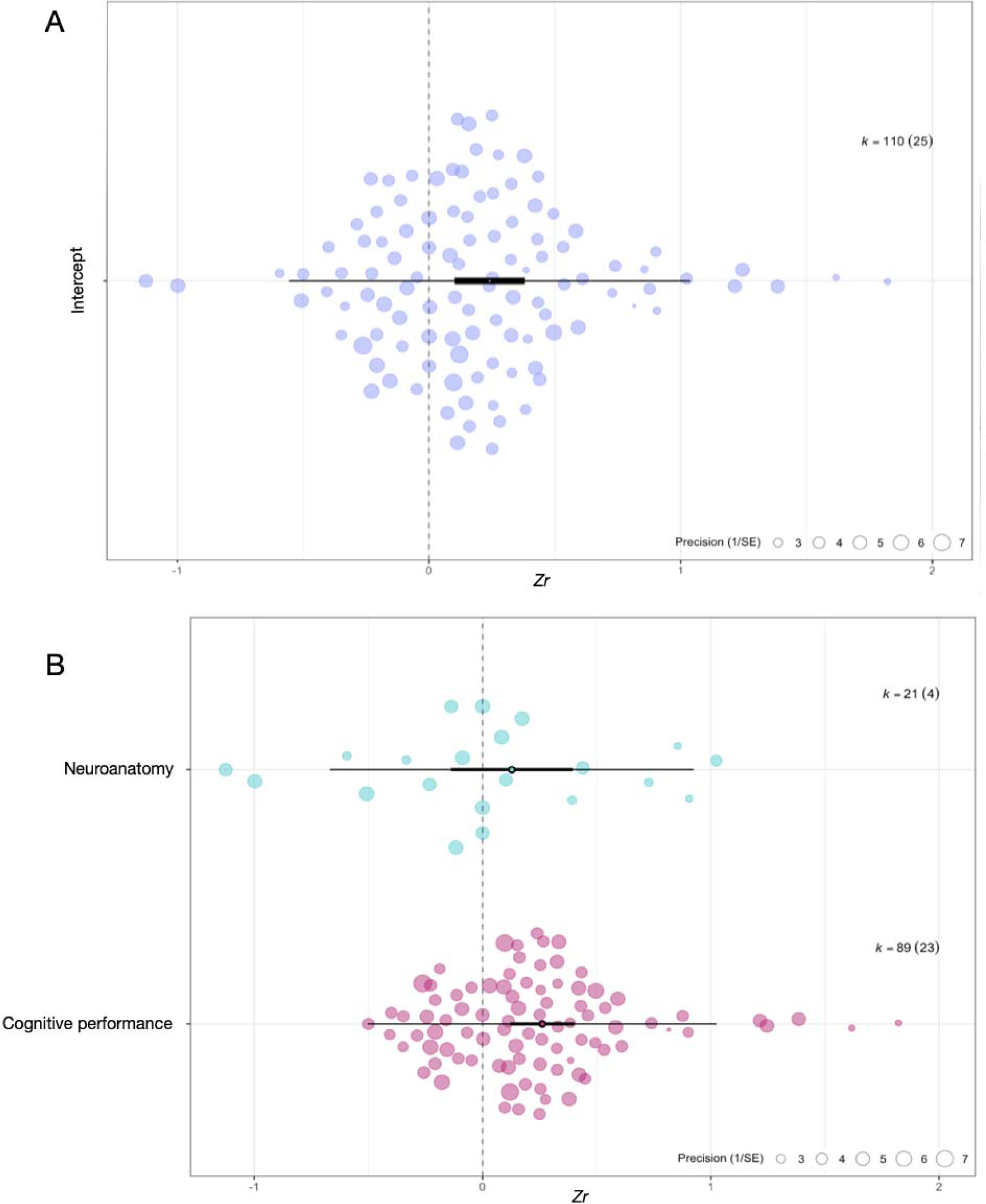
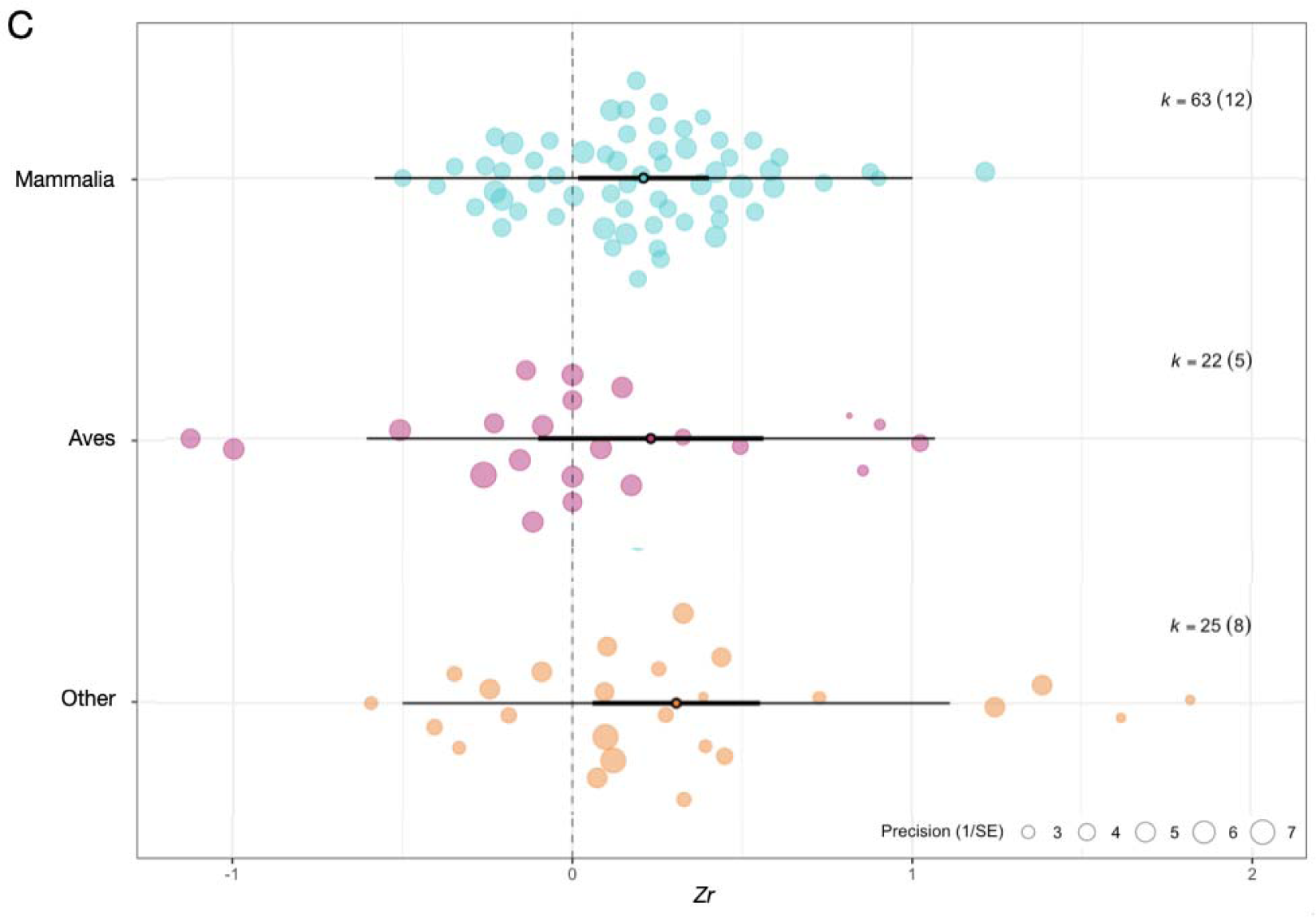
Orchard plots of (A) overall effect size (*Zr*) estimates centred on zero for developmental studies (confidence intervals are positive and did not intersect 0). Orchard plots are also shown for moderators in these studies, including (B) cognition metrics used and (C) taxonomic group. The thick central box represents 95% confidence intervals, the central point is the mean and the thin line represents point estimates. The size of the points represents the precision (1/SE), whereby larger points represent larger sample sizes and thus greater precision.

### (2) Moderator effects

There was high heterogeneity across all four meta-analyses (see Table S5) which motivated further investigation of the factors influencing the relationship between sociality and cognition *via* moderator analyses. However, no moderators explained more variation than the null hypothesis in any of the four meta-analyses (see Figs 4–7 for orchard plots of moderator levels for interspecific studies using individual tests (Fig. 4B–D), interspecific tests using species averages (Fig. 5B–D), intraspecific tests (Fig. 6B–D) and developmental tests (Fig. 7B–D), see also Table S6 for top model sets). Therefore, there is no evidence to suggest a difference in effect size across moderator levels, indicating that the relationship between sociality and cognition is similar across the cognition or sociality metrics used, taxonomic group or testing condition.

### (3) Publication bias

We only detected one type of publication bias in our results. Specifically, developmental studies with smaller sample sizes had larger effect sizes (estimate: 3.23, CI: 0.51/5.95, *P* = 0.02; Fig. S1L). After accounting for this bias, the overall effect size for the relationship between sociality and cognition in developmental studies was substantially reduced and no longer significant (estimate: –0.07, CI: –0.38/0.25).

We did not detect any publication bias according to year of publication in any of the four meta-analyses: interspecific studies based on individual tests – estimate: –0.02, CI: – 0.06/0.02, *P* = 0.41; interspecific studies based on species averages – estimate: –0.12, CI: - 0.04/0.00, *P* = 0.05; intraspecific studies – estimate: 0.01, CI: –0.06/0.09, *P* = 0.66; developmental studies – estimate: –0.01, CI: –0.03/0.01, *P* = 030; Fig. S1). Nor did we detect any other publication bias based on larger effect sizes being published first in the three other meta-analyses (interspecific studies based on individual tests – estimate: 1.21, CI: –0.48/2.90, *P* = 0.15; interspecific studies based on species averages – estimate: -1.65, CI: –4.60/1.29, *P* = 0.26; intraspecific studies – estimate: 1.85, CI: –3.57/7.28, *P* = 0.44, Fig. S1). Given that there is limited evidence of publication bias throughout, this suggests that the results are robust and likely represent a true positive relationship between sociality and cognition.

## IV. DISCUSSION

The SIH was initially conceived as a potential explanation for cognitive evolution in primates (Humphrey, 1976). However, our results reveal that the SIH can be applied as both an evolutionary and developmental explanation of variation in cognitive phenotypes across a range of taxa. We find a significant positive relationship between sociality and cognition across interspecific, intraspecific and developmental studies (however note that we detected potential publication bias in developmental studies), with *r* values ranging from 0.15 to 0.23. Ecological and evolutionary studies tend to have high heterogeneity and thus lower effect sizes because they involve measurements of living animals that are affected by numerous biotic and abiotic factors (Lim, Senior & Nakagawa, 2014). Cohen (1988) initially suggested that effect sizes of less than 0.10 constitute a small effect, and effect sizes of approximately 0.30 constitute a medium effect. However, an assessment of 43 meta-analyses in ecology and evolution has shown that the mean variance explained by *r^2^* was low at 2.51–5.42 and the average effect size in studies using Pearson’s *r* value was 0.18–0.19 (Møller & Jennions, 2002). Additionally, more than 80% of values were smaller than 0.10, with studies in evolutionary biology yielding the lowest values (Møller & Jennions, 2002). Therefore, it was concluded that even minute effect sizes may be important when discussing evolutionary theory (Møller & Jennions, 2002). Consequently, the observed effect size range of 0.15–0.23 that we report here constitutes a biologically meaningful relationship between sociality and cognition and supports the use of the SIH as a leading theory behind the evolution of cognition.

Despite overall support for the SIH, each meta-analysis showed substantial heterogeneity, indicating that additional factors are likely influencing the observed effect sizes, which may in part be because some of the included studies did not directly set out to test the SIH, despite having a social measure which was compared to cognitive performance or neuroanatomy. Therefore, we tested whether several candidate moderator variables explained some of the variation. Our first question was whether evidence for the SIH was influenced by neuroanatomical proxies or behavioural measures of cognition (i.e. cognition tests). Both methods are commonly used to measure cognition, but both have received criticism (Rowe & Healy, 2014; Schubiger, Fichtel & Burkart, 2020; Healy & Rowe, 2007). For example, the relationship between many of the neuroanatomical measures used and cognition is often unclear or debated (Healy & Rowe, 2007). Additionally, many neuroanatomical studies have been criticised for combining data collected using different methods to increase their sample size, as this introduces several confounding factors, such as specimens that are inconsistently measured or preserved (Healy & Rowe, 2007). Likewise, testing cognition behaviourally presents additional challenges as there are many factors that can confound performance, including perceptual and motor bias, motivation, personality, environmental factors, past experience, and heritability [reviewed in Rowe & Healy (2014), Schubiger *et al*. (2020) and Shaw (2017)]. Researchers are also faced with the challenge of devising ways to design tests that compare the same cognitive trait across species, including tasks that account for differences in sensory-motor skills, such as dexterity and colour vision (Shaw & Schmelz, 2017; Schubiger *et al*., 2020; MacLean *et al*., 2014). Ultimately, since effect sizes did not differ significantly between measures of cognition in any of the meta-analyses, the data show that the relationship between sociality and cognition is similar across studies using either neuroanatomy or cognitive performance. Yet despite the lack of statistical significance, there were clear differences in the average effect sizes between cognitive performance and neuroanatomy measures. Effect sizes for cognitive performance were at least twice the size of those for neuroanatomy in three of the meta-analyses (Table 2). Therefore, despite the lack of statistical significance (due to the substantial overlap in variance between the two types of measures), their differences may still be biologically significant.

**Table 2.** Parameter estimates and *P* values for the relationship between sociality and cognition in (A) interspecific studies based on individual tests (*N* = 184 effect sizes from 27 articles); (B) interspecific studies based on species averages (*N* = 129 effect sizes from 36 articles); (C) intraspecific studies (*N* = 161 effect sizes from 16 articles); and (D) developmental studies (*N* = 110 effect sizes from 25 articles). *Zr* is the mean effect size, CI lower and CI upper represent the lower and upper bounds of the 95% confidence interval respectively and *r* is the back-transformed Pearson’s *r* value. See Section II.4 for further details on moderator categories.

|  | Zr | SE | 95%CI<br>Lower | 95%CI<br>Upper | P value | r |
| --- | --- | --- | --- | --- | --- | --- |
| (A) Interspecific (individuals) |  |  |  |  |  |  |
| Meta-analytic mean | 0.16 | 0.10 | −0.03 | 0.35 | 0.09 | 0.16 |
| Phylogenetic mean | 0.16 | 0.10 | −0.04 | 0.36 | 0.09 | 0.16 |
| Cognition metrics |  |  |  |  |  |  |
| Cognitive performance | 0.24 | 0.08 | 0.08 | 0.40 | 0.01 | 0.23 |
| Neuroanatomy | −0.02 | 0.25 | −0.54 | 0.49 | 0.93 | −0.02 |
| Sociality metrics |  |  |  |  |  |  |
| Broad sociality category | 0.26 | 0.11 | 0.05 | 0.48 | 0.02 | 0.26 |
| Group size | −0.06 | 0.15 | −0.35 | 0.23 | 0.68 | −0.06 |
| Social interactions | 0.14 | 0.45 | −0.76 | 1.03 | 0.76 | 0.14 |
| Taxonomy |  |  |  |  |  |  |
| Primates | 0.10 | 0.17 | −0.25 | 0.44 | 0.58 | 0.09 |
| Other | 0.21 | 0.12 | −0.03 | 0.45 | 0.09 | 0.20 |
| (B) Interspecific (species averages) |  |  |  |  |  |  |
| Meta-analytic mean | 0.15 | 0.07 | 0.01 | 0.30 | 0.04 | 0.15 |
| Phylogenetic mean | 0.07 | 0.12 | −0.16 | 0.30 | 0.57 | 0.07 |
| Cognition metrics |  |  |  |  |  |  |
| Cognitive performance | −0.04 | 0.09 | −0.23 | 0.15 | 0.67 | −0.04 |
| Neuroanatomy | 0.15 | 0.08 | −0.01 | 0.31 | 0.07 | 0.15 |
| Sociality metrics |  |  |  |  |  |  |
| Broad sociality category | 0.08 | 0.13 | −0.19 | 0.35 | 0.56 | 0.08 |
| Group size | 0.19 | 0.07 | 0.04 | 0.34 | 0.01 | 0.19 |
| Social interactions | 0.29 | 0.28 | −0.27 | 0.85 | 0.30 | 0.28 |
| Taxonomy |  |  |  |  |  |  |
| Primates | 0.25 | 0.10 | 0.05 | 0.45 | 0.02 | 0.25 |
| Other | 0.05 | 0.09 | −0.13 | 0.24 | 0.55 | 0.05 |
| (C) Intraspecific |  |  |  |  |  |  |
| Meta-analytic mean | 0.23 | 0.10 | 0.04 | 0.42 | 0.02 | 0.23 |
| Phylogenetic mean | 0.23 | 0.10 | 0.04 | 0.42 | 0.02 | 0.22 |
| Cognition metrics |  |  |  |  |  |  |
| Cognitive performance | 0.44 | 0.13 | 0.18 | 0.70 | <0.001 | 0.41 |
| Neuroanatomy | 0.10 | 0.08 | −0.07 | 0.26 | 0.24 | 0.10 |
| Sociality metrics |  |  |  |  |  |  |
| Broad sociality category | 0.01 | 0.14 | −0.29 | 0.31 | 0.94 | 0.01 |
| Group size | 0.33 | 0.11 | 0.08 | 0.58 | 0.01 | 0.32 |
| Social interactions | 0.31 | 0.21 | −0.14 | 0.75 | 0.16 | 0.30 |
| **Testing conditions** |  |  |  |  |  |  |
| Captive | 0.28 | 0.11 | 0.05 | 0.50 | 0.02 | 0.27 |
| Wild | 0.24 | 0.16 | −0.12 | 0.59 | 0.18 | 0.23 |
| Unknown | −0.23 | 0.03 | −0.28 | −0.17 | <0.001 | −0.22 |
| **(D) Developmental** |  |  |  |  |  |  |
| Meta-analytic mean | 0.24 | 0.07 | 0.11 | 0.38 | <0.01 | 0.24 |
| Phylogenetic mean | 0.24 | 0.07 | 0.11 | 0.38 | <0.01 | 0.24 |
| **Cognition metrics** |  |  |  |  |  |  |
| Cognitive performance | 0.26 | 0.07 | 0.13 | 0.40 | <0.001 | 0.26 |
| Neuroanatomy | 0.13 | 0.17 | −0.22 | 0.48 | 0.46 | 0.13 |
| **Taxonomy** |  |  |  |  |  |  |
| Mammals | 0.21 | 0.06 | 0.08 | 0.34 | <0.01 | 0.21 |
| Birds | 0.21 | 0.25 | −0.29 | 0.75 | 0.36 | 0.23 |
| Other | 0.31 | 0.15 | 0.00 | 0.48 | 0.61 | 0.30 |

Although effect sizes did not differ significantly between measures of sociality (indicating that support for the SIH is not predicted by the measure of sociality used), there was an overwhelming lack of diversity in the social measures used to test the SIH. Given that the SIH is based on the premise of cognitive challenges, such as needing to recognise and respond to a variety of social situations, tests of a relationship between sociality and cognition rarely measure multiple components and dimensions of sociality beyond social group size. Several innovative techniques have been developed to quantify social complexity which may provide further insight into the SIH. For instance, Bergman & Beehner (2015) proposed that social complexity could be measured as the number of differentiated relationships within a group, as distinguishing amongst individuals requires cognitive traits such as discrimination and memory. Other studies have also proposed alternative methods for examining social complexity based on association indices that reflect different types of dyadic relationships within a network (Bergman & Beehner, 2015; Fischer *et al*., 2017). Thus, we encourage future studies to explore these alternative methods of quantifying the social environment, and in turn, investigate their relationship with cognition.

Although the SIH was initially formulated to explain primate cognitive evolution, evidence from multiple taxonomic groups suggests that the SIH can be applied to a much broader range of species than initially hypothesised. The low heterogeneity value for species identity across developmental and interspecific studies (Table S5) suggests that there was no significant phylogenetic signal, with similar effect sizes across the taxonomic groups tested, despite the relatively poor taxonomic representation available. Due to a lack of taxonomic diversity, we were only able to investigate taxonomy at the class or order level. We did not detect an effect of taxonomy at either class or order level (although some results between primates and non-primates were borderline, particularly in interspecific studies). Although the SIH has expanded beyond primates, studies are largely restricted to a handful of species or taxonomic groups. Tests of the SIH have been dominated by mammals, specifically primates and rodents, as well as birds, specifically passerines. For example, 29% of studies on rodents come from *Mus musculus* (Fitchett, Barnard & Cassaday, 2009; Kogan *et al*., 2000; Smith *et al*., 2018; Wang *et al*., 2018; Williams *et al*., 2001) and a further 36% come from *Rattus norvegicus* (Toyoshima *et al*., 2018; Schrijver *et al*., 2002; Heimer-McGinn *et al*., 2020; Dalley *et al*., 2002; Brenes *et al*., 2016). There was also a noticeable lack of studies on amphibians and reptiles. Of the three studies on reptiles, (Riley *et al*., 2018, 2017; De Meester *et al*., 2019), two were on tree skinks (*Egernia stiolata*) and revealed no support for the SIH (Riley *et al*., 2018, 2017). Additionally, there was a surprising lack of diversity in the species of insects tested given the vast size of the order, with tests dominated by bees, wasps, ants and locusts (Hymenoptera and Orthoptera, Fig. 3) (Kamhi *et al*., 2016; O’Donnell *et al*., 2019, 2015; Ott & Rogers, 2010; Rehan, Bulova & O’Donnell, 2015; Smith *et al*., 2010; Seid & Junge, 2016; Amador-Vargas *et al*., 2015). We encourage future studies to expand tests to a broader diversity of species, as this will be crucial to determining further the generalisability of the SIH across taxonomic groups.

Finally, testing conditions did not impact the observed effect sizes, with similar estimates produced for captive and wild studies. These results suggest that support for the SIH is robust across testing conditions and reinforces that both situations are valid for testing the SIH. It also reinforces the idea that studies of captive animals may be useful in providing controlled conditions for experimental manipulations that can assist in teasing apart causal relationships and disentangling specific factors affecting cognitive processes (Szabo *et al*., 2022). By contrast, studies of free-living animals may be useful to link cognitive traits to their function by testing in ecologically relevant conditions (Harrison & van de Waal, 2022). It was notable that no SIH studies adopted a matched-species comparison (De Petrillo, Bettle & Rosati, 2022), testing individuals in both captivity and the wild. Ultimately, to obtain a more comprehensive picture of the factors affecting cognitive processes and to link cognitive abilities to their function, we advocate that future studies implement a matched-species comparison.

There was limited evidence of publication bias, apart from in developmental studies. This suggests that our results for inter- and intraspecific meta-analyses are robust. However, developmental studies showed a bias toward smaller sample sizes yielding larger effect sizes, which could leverage the results in favour of the SIH. This bias, termed small-study effects, may be due to studies with smaller sample sizes being more vulnerable to the effects of heterogeneity (Sterne, Gavaghan & Egger, 2000). Additionally, studies on juveniles are often prone to smaller sample sizes as individuals may be lost to illness or predation during longitudinal testing. After applying a correction for the bias, the effect size of development studies was substantially reduced which suggests that these results are leveraging the data set, because when the sample size is infinitely large the estimated effect size becomes smaller and confidence intervals intersect zero. However, this correction is an extrapolation and may not reflect the true effect size. Therefore, both interpretations (an effect of development versus an effect due to publication bias) should be treated with caution.

Initially, we intended to investigate cognitive metrics in greater detail. Specifically, we wanted to separate neuroanatomical measures into brain regions to determine whether specific regions were associated with greater support for the SIH. Likewise, we intended to break down cognitive tests into domain-general and socio-cognitive tests to examine whether support for the hypothesis extended beyond social cognition. However, the unequal distribution of brain regions [studies of absolute/relative brain size (e.g. Shultz & Dunbar, 2010; MacLean *et al*., 2009; Dunbar & Shultz, 2007; Kverková *et al*., 2018; Weisbecker *et al*., 2015) and neocortex (Schillaci, 2008; Sandel *et al*., 2016) vastly outnumbered studies on other brain regions] prevented us from analysing this relationship at a finer scale than comparing cognitive tests to neuroanatomy. Additionally, given that the SIH is based on the cognitive demands of sociality, it was surprising to see how few studies utilise social tests of cognition such as tactical deception (Byrne & Corp, 2004), theory of mind (Devaine *et al*., 2017), social learning (Sewall *et al*., 2018; Lefebvre, Palameta & Hatch, 1996) and inequity aversion (Wascher, 2015). Many intraspecific tests of the relationship between sociality and cognition use domain-general tasks, rather than socio-cognitive tasks to measure cognition, despite the explicit prediction that *social* pressures have driven the evolution of cognition (Dunbar, 1998; Jolly, 1966; Humphrey, 1976; Chance & Mead, 1953). Continuing to expand the cognitive metrics tested both behaviourally and neuroanatomically, especially expanding research on socio-cognitive metrics, represents a promising avenue for future research.

## V. CONCLUSIONS

(1) Overall, we detected support for the SIH in the literature across a wide range of studies, highlighting the importance of this hypothesis as a core theory behind our understanding of the evolution of cognition and cognitive development.

(2) Effect sizes did not differ significantly between the cognitive or sociality metrics used, taxonomy, or testing conditions, which suggests that support for the SIH is similar across studies using neuroanatomy and cognitive performance, those using broad categories of sociality, group size and social interactions, among taxonomic groups, and for tests conducted in captivity or the wild. This suggests that the SIH is more broadly applicable than originally hypothesised.

(3) Since many taxa are underrepresented, we encourage an expansion of the taxonomic groups tested in future studies of the SIH, as well as the development of standardised cognitive test batteries to facilitate cross-species comparisons of cognitive performance.

(4) We also encourage the inclusion of social cognition tests, as well as standardised neuroanatomical testing methods, to clarify whether the SIH predominantly applies to social cognition, or domain-general cognition.

(5) To facilitate future analysis of the SIH we advocate for increased utilisation of innovative programmes such as ManyPrimates (Atschul *et al*., 2019) and ManyBirds (Lambert *et al*., 2022) that allow researchers to collaborate, jointly discuss ideas and develop standardised testing procedures. These programmes aim to give further insight into the evolution of primate and avian cognition by pooling data from multiple laboratories and thus increase species representation and sample sizes across studies.

## VI. ACKNOWLEDGEMENTS

We thank Phoebe Ranford for her assistance with assessing whether 10% of papers met inclusion criteria, Canaan Perry for his expertise and advice with setting up the review search and Camilla Soravia for her feedback on manuscript drafts. We also thank the University of Western Australia for logistical support, the Australian government for the Research Training Stipend scholarship support to E.M.S. and the Australian Research Council funding to A.R.R. and B.J.A.

## VII. DATA ACCESSIBILITY

Data and code can be accessed at 10.6084/m9.figshare.25533115.

## IX. SUPPORTING INFORMATION

Additional supporting information may be found online in the Supporting Information section at the end of the article.

**Table S1.** List of all papers included in all four meta-analyses.

**Table S2.** Comparison of null model and a model with taxonomic group as a predictor in intraspecific studies.

**Table S3.** Comparison of null model and a model with testing conditions as a predictor in intraspecific studies.

**Table S4.** *R*^2^ values (proportion of variance) explained by each moderator in each meta-analysis.

**Table S5.** Heterogeneity (*I*^2^) values for each of the random effects in each meta-analysis.

**Table S6.** Top model set of candidate terms affecting the correlation between sociality and cognition for each study category.

**Fig. S1.** Publication bias tests for interspecific studies using individual tests, interspecific studies using species averages, intraspecific studies and developmental studies.

## Notes

https://doi.org/10.6084/m9.figshare.25533115.v1

### Competing Interest Statement

The authors have declared no competing interest.

## VIII. REFERENCES

Amador-Vargas, S., Gronenberg, W., Wcislo, W. T. & Mueller, U. (2015). Specialization and group size: Brain and behavioural correlates of colony size in ants lacking morphological castes. Proceedings of the Royal Society B: Biological Sciences 282(1801), 20142502.

American Psychological Association (2022). Sociality. APA dictionary of psychology.

Amici, F., Aureli, F. & Call, J. (2008). Fission-fusion dynamics, behavioral flexibility, and inhibitory control in primates. Current Biology 18(18), 1415–1419.

Aplin, L. M., Farine, D. R., Morand-Ferron, J. & Sheldon, B. C. (2012). Social networks predict patch discovery in a wild population of songbirds. Proceedings of the Royal Society B: Biological Sciences 279(1745), 4199–4205.

Arsznov, B. M. & Sakai, S. T. (2012). Pride diaries: Sex, brain size and sociality in the African lion (*Panthera leo*) and cougar (*Puma concolor*). Brain, Behavior and Evolution 79(4), 275–289.

Arsznov, B. M. & Sakai, S. T. (2013). The Procyonid social club: Comparison of brain volumes in the coatimundi (*Nasua nasua, N. narica*), Kinkajou (*Potos flavus*), and Raccoon (*Procyon lotor*). Brain, Behavior and Evolution 82(2), 129–145.

Ash, H., Ziegler, T. E. & Colman, R. J. (2020). Early learning in the common marmoset *(Callithrix jacchus*): Behavior in the family group is related to preadolescent cognitive performance. American Journal of Primatology 82(8), e23159.

Ashton, B. J., Ridley, A. R., Edwards, E. K. & Thornton, A. (2018a). Cognitive performance is linked to group size and affects fitness in Australian magpies. Nature 554, 364–367.

Ashton, B. J., Thornton, A. & Ridley, A. R. (2018b). An intraspecific appraisal of the social intelligence hypothesis. Philosophical Transactions of the Royal Society B: Biological Sciences 373, 20170288.

Ashton, B. J., Thornton, A. & Ridley, A. R. (2019). Larger group sizes facilitate the emergence and spread of innovations in a group-living bird. Animal Behaviour 158, 1–7.

Atschul, D. M., Berhan, M. J., Bohn, M., Caspar, K. R., Fitchel, C., Försterling, M., Grebe, N. M., Hernandez-Aguilar, A. R., Kwok, S. C., Llorente, M., Motes-Rorigo, A., Proctor, D., SàNchez-Amaro, A., Simpson, E. A., Szabelska, A., et al. (2019). Collaborative open science as a way to reproducibility and new insights in primate cognition research. Japanese Psychological Review 62(3), 205–220.

Ausas, M. S., Mazzitelli-Fuentes, L., Roman, F. R., Crichigno, S. A., DE Vincenti, A. P. & Mongiat, L. A. (2019). Social isolation impairs active avoidance performance and decreases neurogenesis in the dorsomedial telencephalon of rainbow trout. Physiology and Behavior 198, 1–10.

Bannier, F., Tebbich, S. & Taborsky, B. (2017). Early experience affects learning performance and neophobia in a cooperatively breeding cichlid. Ethology 123(10), 712–723.

Bartoń, K. (2022). MuMIn: Multi-model inference, vol. R package version 1.471.1, https://cran.r-project.org/package=MuMIn.

Barton, R. A. (1996). Neocortex size and behavioural ecology in primates. Proceedings of the Royal Society B: Biological Sciences 263(1367), 173–177.

Beauchamp, G. & FERNÁNDEZ-Juricic, E. (2004). Is there a relationship between forebrain size and group size in birds? Evolutionary Ecology Research 6(6), 833–842.

Bergman, T. J. & Beehner, J. C. (2015). Measuring social complexity. Animal Behaviour 103, 203–209.

Berhane, J. F. & Gazes, R. P. (2020). Social monkeys learn more slowly: Social network centrality and age are positively related to learning errors by capuchin monkeys (*Cebus [Sapajus] apella*). Canadian Journal of Experimental Psychology 74(3), 228– 234.

Bond, A. B., Kamil, A. C. & Balda, R. P. (2003). Social complexity and transitive inference in corvids. Animal Behaviour 65(3), 479–487.

Bond, A. B., Kamil, A. C. & Balda, R. P. (2007). Serial reversal learning and the evolution of behavioral flexibility in three species of North American corvids (*Gymnorhinus cyanocephalus, Nucifraga columbiana*, Aphelocoma californica). Journal of Comparative Psychology 121(4), 372–379.

Bond, A. B., Wei, C. A. & Kamil, A. C. (2010). Cognitive representation in transitive inference: A comparison of four corvid species. Behavioural Processes 85(3), 283– 292.

Boogert, N. J., Farine, D. R. & Spencer, K. A. (2014). Developmental stress predicts social network position. Biology Letters 10, 20140561.

Boogert, N. J., Madden, J. R., MORAND-Ferron, J. & Thornton, A. (2018). Measuring and understanding individual differences in cognition. Philosophical Transactions of the Royal Society B: Biological Sciences 373, 20170280.

Borrego, N. & Gaines, M. (2016). Social carnivores outperform asocial carnivores on an innovative problem. Animal Behaviour 114, 21–26.

Brandão, M. L., Braithwaite, V. A. & GONCALVES-DE-Freitas, E. (2015). Isolation impairs cognition in a social fish. Applied Animal Behaviour Science 171, 204–210. Brenes, J. C., Lackinger, M., Höglinger, G. U., Schratt, G., Schwarting, R. K. W. &

Wöhr, M. (2016). Differential effects of social and physical environmental enrichment on brain plasticity, cognition, and ultrasonic communication in rats. Journal of Comparative Neurology 524(8), 1586–1607.

Burish, M. J., Kueh, H. Y. & Wang, S. S. H. (2004). Brain architecture and social complexity in modern and ancient birds. Brain Behavior and Evolution 63(2), 107– 124.

Byrne, R. W. & Corp, N. (2004). Neocortex size predicts deception rate in primates. Proceedings of the Royal Society B: Biological Sciences 271(1549), 1693–1699.

Byrne, R. W. & Whiten, A. (1988). Machiavellian intelligence: Social expertise and the evolution of intellect in monkeys, apes, and humans. Clarendon Press, Oxford.

Carducci, J. P. & Jakob, E. M. (2000). Rearing environment affects behaviour of jumping spiders. Animal Behaviour 59, 39–46.

Cazakoff, B. N., Johnson, K. J. & Howland, J. G. (2010). Converging effects of acute stress on spatial and recognition memory in rodents: A review of recent behavioural and pharmacological findings. Progress in Neuro-Psychopharmacology & Biological Psychiatry 34(5), 733–741.

Chance, M. & Mead, A. (1953). Social behaviour and primate evolution. Symposia of the Society for Experimental Biology 7, 395–439.

Cohen, J. (1988). Statistical power analysis for the behavioral sciences, 2nd edition. Routledge, New York.

Costanzo, M. S., Bennett, N. C. & Lutermann, H. (2009). Spatial learning and memory in African mole-rats: The role of sociality and sex. Physiology and Behavior 96, 128– 134.

Cowl, V. B. & Shultz, S. (2017). Large brains and groups associated with high rates of agonism in primates. Behavioral Ecology 28(3), 803–810.

Dalley, J., Theobald, D., Pereira, E., Li, P. & Robbins, T. (2002). Specific abnormalities in serotonin release in the prefrontal cortex of isolation-reared rats measured during behavioural performance of a task assessing visuospatial attention and impulsivity. Psychopharmacology 164(3), 329–340.

Darwin, C. (1871). The descent of man and selection in relation to sex. John Murray Press, London.

DE Meester, G., Huyghe, K. & Van Damme, R. (2019). Brain size, ecology and sociality: A reptilian perspective. Biological Journal of the Linnean Society 126(3), 381–391.

DE Petrillo, F., Bettle, R. & Rosati, A. G. (2022). Insights from matched species comparisons for understanding cognition in the wild. Current Opinion in Behavioral Sciences 45, 101134.

Dean, L. G., Hoppitt, W., Laland, K. N. & Kendal, R. L. (2011). Sex ratio affects sex-specific innovation and learning in captive ruffed lemurs (*Varecia variegata* and *Varecia rubra*). American Journal of Primatology 73(12), 1210–1221.

Deaner, R. O., Nunn, C. L. & VAN Schaik, C. P. (2000). Comparative tests of primate cognition: Different scaling methods produce different results. Brain Behavior and Evolution 55, 44–52.

Decasien, A. R. & Higham, J. P. (2019). Primate mosaic brain evolution reflects selection on sensory and cognitive specialization. Nature Ecology & Evolution 3(10), 1483– 1493.

Decasien, A. R., Williams, S. A. & Higham, J. P. (2017). Primate brain size is predicted by diet but not sociality. Nature Ecology & Evolution 1(5), 0112.

Devaine, M., San-Galli, A., Trapanese, C., Bardino, G., Hano, C., SAINT Jalme, M., Bouret, S., Masi, S. & Daunizeau, J. (2017). Reading wild minds: A computational assay of theory of mind sophistication across seven primate species. PLoS Computational Biology 13(11), e1005833.

Dunbar, R. I. M. (1992). Neocortex size as a constraint on group size in primates. Journal of Human Evolution 22(6), 469–493.

Dunbar, R. I. M. (1995). Neocortex size and group size in primates: A test of the hypothesis. Journal of Human Evolution 28(3), 287–296.

Dunbar, R. I. M. (1998). The social brain hypothesis. *Evolutionary Anthropology: Issues*, News, and Reviews 6, 178–190.

Dunbar, R. I. M. & Bever, J. (1998). Neocortex size predicts group size in carnivores and some insectivores. Ethology 104(8), 695–708.

Dunbar, R. I. M. & Shultz, S. (2007). Understanding primate brain evolution. Philosophical Transactions of the Royal Society B: Biological Sciences 362(1480), 649–658.

Fedorova, N., Evans, C. L. & Byrne, R. W. (2017). Living in stable social groups is associated with reduced brain size in woodpeckers (Picidae). Biology Letters 13(3), 20170008.

Fichtel, C., Dinter, K. & Kappeler, P. M. (2020). The lemur baseline: How lemurs compare to monkeys and apes in the primate cognition test battery. PeerJ 8, e10025.

Fischer, J., Farnworth, M. S., SENNHENN-Reulen, H. & Hammerschmidt, K. (2017). Quantifying social complexity. Animal Behaviour 130, 57–66.

Fischer, S., BESSERT-Nettelbeck, M., Kotrschal, A. & Taborsky, B. (2015). Rearing-group size determines social competence and brain structure in a cooperatively breeding cichlid. The American Naturalist 186, 123–140.

Fitchett, A. E., Barnard, C. J. & Cassaday, H. J. (2009). Corticosterone differences rather than social housing predict performance of T-maze alternation in male CD-I mice. Animal Welfare 18, 21–31.

Forss, S. I. F., Willems, E., Call, J. & VAN Schaik, C. P. (2016). Cognitive differences between orang-utan species: A test of the cultural intelligence hypothesis. Scientific Reports 6, 30516.

Fox, J. & Monette, G. (1992). Generalized collinearity diagnostics. Journal of the American Statistical Association 87(417), 178–183.

Fox, K. C. R., Muthukrishna, M. & Shultz, S. (2017). The social and cultural roots of whale and dolphin brains. Nature Ecology & Evolution 1, 1699–1705.

Freeberg, T. M., Dunbar, R. I. M. & Ord, T. J. (2012). Social complexity as a proximate and ultimate factor in communicative complexity. Philosophical Transactions of the Royal Society B: Biological Sciences 367, 1785–1801.

Gil, D., Naguib, M., Riebel, K., Rutstein, A. & Gahr, M. (2006). Early condition, song learning, and the volume of song brain nuclei in the zebra finch (*Taeniopygia guttata*). Journal of Neurobiology 66(14), 1602–1612.

Graham, K. L. (2011). Coevolutionary relationship between striatum size and social play in nonhuman primates. American Journal of Primatology 73(4), 314–322.

Grueber, C. E., Nakagawa, S., Laws, R. J. & Jamieson, I. G. (2011). Multimodel inference in ecology and evolution: Challenges and solutions. Journal of Evolutionary Biology 24(4), 699–711.

Güntürkün, O. & Bugnyar, T. (2016). Cognition without cortex. Trends in Cognitive Sciences 20(4), 291–303.

HARRELL JR., F. E., Lee, K. L. & Mark, D. B. (1996). Multivariate prognostic models: Issues in developing models, evaluating assumptions and adequacy, and measuring and reducing errors. Statistics in Medicine 15(4), 361–387.

Harris, A. P., D’Eath, R. B. & Healy, S. D. (2009). Environmental enrichment enhances spatial cognition in rats by reducing thigmotaxis (wall hugging) during testing. Animal Behaviour 77(6), 1459–1464.

Harrison, R. A. & VAN DE Waal, E. (2022). The unique potential of field research to understand primate social learning and cognition. Current Opinion in Behavioral Sciences 45, 101132.

Healy, S. D. & Rowe, C. (2007). A critique of comparative studies of brain size. Proceedings of the Royal Society B: Biological Sciences 274, 453–464.

Healy, S. D. & Rowe, C. (2013). Costs and benefits of evolving a larger brain: doubts over the evidence that large brains lead to better cognition. Animal Behaviour 86, e1–e3.

Hedges, L. & Olkin, I. (1985). Statistical methods for meta-analysis. Academic Press, New York.

Heimer-Mcginn, V. R., Wise, T. B., Hemmer, B. M., Dayaw, J. N. T. & Templer, V. L. (2020). Social housing enhances acquisition of task set independently of environmental enrichment: A longitudinal study in the Barnes maze. Learning & Behavior 48(3), 322–334.

Higgins, J. P. & Thompson, S. G. (2002). Quantifying heterogeneity in a meta-analysis. Statistics in Medicine 21(11), 1539–1558.

Higgins, J. P. T., Thompson, S. G., Deeks, J. J. & Altman, D. G. (2003). Measuring inconsistency in meta-analyses. BMJ 327(7414), 557–560.

Hinchliff, C. E., Smith, S. A., Allman, J. F., Burleigh, J. G., Chaudhary, R., Coghill, L. M., Crandall, K. A., Deng, J., Drew, B. T., Gazis, R., Gude, K., Hibbett, D. S., Katz, L. A., Laughinghouse, H. D., Mctavish, E. J., et al. (2015). Synthesis of phylogeny and taxonomy into a comprehensive tree of life. Proceedings of the National Academy of Sciences 112(41), 12764–12769.

Hobson, E. A., Ferdinand, V., Kolchinsky, A. & Garland, J. (2019). Rethinking animal social complexity measures with the help of complex systems concepts. Animal Behaviour 155, 287–296.

Humphrey, N. K. (1976). The social function of intellect. In Growing points in ethology (ed. P. P. G. Bateson and R. A. Hinde), pp. 303–317. Cambridge University Press, Oxford, England.

Iwaniuk, A. N. & Arnold, K. E. (2004). Is cooperative breeding associated with bigger brains? A comparative test in the Corvida (Passeriformes). Ethology 110(3), 203–220.

Jennions, M. D. & Møller, A. P. (2002). Relationships fade with time: A meta-analysis of temporal trends in publication in ecology and evolution. Proceedings of the Royal Society B: Biological Sciences 269(1486), 43–48.

Jerison, H. J. (1973). Evolution of the brain and intelligence. Academic Press, New York.

Joffe, T. H. & Dunbar, R. I. M. (1997). Visual and socio-cognitive information processing in primate brain evolution. Proceedings of the Royal Society B: Biological Sciences 264(1386), 1303–1307.

Johnson-Ulrich, L. & Holekamp, K. E. (2020). Group size and social rank predict inhibitory control in spotted hyaenas. Animal Behaviour 160, 157–168.

Jolly, A. (1966). Lemur social behavior and primate intelligence. Science 153, 501–506.

Kamhi, J. F., Gronenberg, W., Robson, S. K. A. & Traniello, J. F. A. (2016). Social complexity influences brain investment and neural operation costs in ants. Proceedings of the Royal Society B: Biological Sciences 283(1841), 20161949.

Kelly, E. M. (2016). Counting on your friends: The role of social environment on quantity discrimination. Behavioural Processes 128, 9–16.

Keverne, E. B., Martel, F. L. & Nevison, C. M. (1996). Primate brain evolution: Genetic and functional considerations. Proceedings of the Royal Society B: Biological Sciences 263(1371), 689–696.

Kogan, J. H., Frankland, P. W. & Silva, A. J. (2000). Long-term memory underlying hippocampus-dependent social recognition in mice. Hippocampus 10, 47–56.

Krasheninnikova, A., Brager, S. & Wanker, R. (2013). Means-end comprehension in four parrot species: Explained by social complexity. Animal Cognition 16(5), 755– 764.

Kudo, H. & Dunbar, R. I. M. (2001). Neocortex size and social network size in primates. Animal Behaviour 62, 711–722.

Kulahci, I. G., Ghazanfar, A. A. & Rubenstein, D. I. (2018). Knowledgeable lemurs become more central in social networks. Current Biology 28, 1306–1310.

Kulahci, I. G., Rubenstein, D. I., Bugnyar, T., Hoppitt, W., Mikus, N. & Schwab, C. (2016). Social networks predict selective observation and information spread in ravens. Royal Society Open Science 3(7), 160256.

Kverková, K., Belikova, T., Olkowicz, S., Pavelkova, Z., O’riain, M. J., Sumbera, R., Burda, H., Bennett, N. C. & Nemec, P. (2018). Sociality does not drive the evolution of large brains in eusocial African mole-rats. Scientific Reports 8, 9203– 9214.

Lambert, M., Farrar, B., Garcia-Pelegrin, E., Reber, S. A. & Miller, R. (2022). ManyBirds: A multi-site collaborative Open Science approach to avian cognition and behaviour research. Animal Behaviour and Cognition 9, 133–152.

Langley, E. J. G., VAN Horik, J. O., Whiteside, M. A., Beardsworth, C. E. & Madden, J. R. (2018a). The relationship between social rank and spatial learning in pheasants, *Phasianus colchicus*: Cause or consequence? PeerJ 6, e5738.

Langley, E. J. G., Van Horik, J. O., Whiteside, M. A. & Madden, J. R. (2018b). Individuals in larger groups are more successful on spatial discrimination tasks. Animal Behaviour 142, 87–93.

Lefebvre, L., Palameta, B. & Hatch, K. K. (1996). Is group-living associated with social learning? A comparative test of a gregarious and a territorial columbid. Behaviour 133, 241–261.

Lehmann, J. & Dunbar, R. I. M. (2009). Network cohesion, group size and neocortex size in female-bonded Old World primates. Proceedings of the Royal Society B: Biological Sciences 276(1677), 4417–4422.

Leris, I. & Reader, S. M. (2016). Age and early social environment influence guppy social learning propensities. Animal Behaviour 120, 11–19.

Lewis, K. P. & Barton, R. A. (2006). Amygdala size and hypothalamus size predict social play frequency in nonhuman primates: A comparative analysis using independent contrasts. Journal of Comparative Psychology 120, 31–37.

Liberati, A., Altman, D. G., Tetzlaff, J., Mulrow, C., Gøtzsche, P. C., Ioannidis, J. P., Clarke, M., Devereaux, P. J., Kleijnen, J. & Moher, D. (2009). The PRISMA statement for reporting systematic reviews and meta-analyses of studies that evaluate healthcare interventions: Explanation and elaboration. BMJ 339, b2700.

Liedtke, J. & Schneider, J. M. (2017). Social makes smart: Rearing conditions affect learning and social behaviour in jumping spiders. Animal Cognition 20, 1093–1106.

Liker, A. & Bókony, V. (2009). Larger groups are more successful in innovative problem solving in house sparrows. Proceedings of the National Academy of Sciences 106(19), 7893–7898.

Lim, J. N., Senior, A. M. & Nakagawa, S. (2014). Heterogeneity in individual quality and reproductive trade-offs within species. Evolution 68(8), 2306–2318.

Lindenfors, P. (2005). Neocortex evolution in primates: The ‘social brain’ is for females. Biology Letters 1(4), 407–410.

Lindenfors, P., Nunn, C. L. & Barton, R. A. (2007). Primate brain architecture and selection in relation to sex. BMC Biology 5, 20.

Lipkind, D., Nottebohm, F., Rado, R. & Barnea, A. (2002). Social change affects the survival of new neurons in the forebrain of adult songbirds. Behavioural Brain Research 133, 31–43.

Lipsey, M. W. & Wilson, D. B. (2001). Practical meta-analysis. Sage Publications, Inc, Thousand Oaks, California.

Logan, C. J., Jelbert, S., Lukas, D., Buskell, A., Navarrete, A. F., Cross, F. R., CURRIE, A. & Montgomery, S. H. (2018). Beyond brain size: Uncovering the neural correlates of behavioral and cognitive specialization. Comparative Cognition and Behavior Reviews 13, 55–89.

Louail, M., Gilissen, E., Prat, S., Garcia, C. & Bouret, S. (2019). Refining the ecological brain: Strong relation between the ventromedial prefrontal cortex and feeding ecology in five primate species. Cortex 118, 262–274.

Machatschke, I. H., Bauer, B., Glenk, L. M., Millesi, E. & Wallner, B. (2011). Spatial learning and memory differs between single and cohabitated guinea pigs. Physiology and Behavior 102, 311–316.

Maclean, E. L., Barrickman, N. L., Johnson, E. M. & Wall, C. E. (2009). Sociality, ecology, and relative brain size in lemurs. Journal of Human Evolution 56(5), 471– 478.

Maclean, E. L., Hare, B., Nunn, C. L., Addessi, E., Amici, F., Anderson, R. C., Aureli, F., Baker, J. M., Bania, A. E., Barnard, A. M., Boogert, N. J., Brannon, E. M., Bray, E. E., Bray, J., Brent, L. J., et al. (2014). The evolution of self-control. Proceedings of the National Academy of Sciences 111(20), e2140–e2148.

Maclean, E. L., Merritt, D. J. & Brannon, E. M. (2008). Social complexity predicts transitive reasoning in prosimian primates. Animal Behaviour 76, 479–486.

Maclean, E. L., Sandel, A. A., Bray, J., Oldenkamp, R. E., Reddy, R. B. & Hare, B. A. (2013). Group size predicts social but not nonsocial cognition in lemurs. PLOS One 8, e66359.

Meagher, R. K., Daros, R. R., Costa, J. H. C., VON Keyserlingk, M. A. G., Hötzel, M. J. & Weary, D. M. (2015). Effects of degree and timing of social housing on reversal learning and response to novel objects in dairy calves. PLOS One 10(8), e0132828.

Meguerditchian, A., Marie, D., Margiotoudi, K., Roth, M., Nazarian, B., Anton, J. L. & Claidiere, N. (2021). Baboons (*Papio anubis*) living in larger social groups have bigger brains. Evolution and Human Behavior 42, 30–34.

Michonneau, F., Brown, J. W. & Winter, D. J. (2016). rotl: An R package to interact with the Open Tree of Life data. Methods in Ecology and Evolution 7(12), 1476–1481.

Morand-Ferron, J., Cole, E. F. & Quinn, J. L. (2016). Studying the evolutionary ecology of cognition in the wild: A review of practical and conceptual challenges. Biological Reviews 91, 367–389.

Morand-Ferron, J. & Quinn, J. L. (2011). Larger groups of passerines are more efficient problem solvers in the wild. Proceedings of the National Academy of Sciences 108(38), 15898–15903.

Morrison, R. E., Eckardt, W., Stoinski, T. S. & Brent, L. J. N. (2020). Comparing measures of social complexity: Larger mountain gorilla groups do not have a greater diversity of relationships. Proceedings of the Royal Society B: Biological Sciences 287(1931), 20201026.

Møller, A. & Jennions, M. D. (2002). How much variance can be explained by ecologists and evolutionary biologists? Oecologia 132(4), 492–500.

Nakagawa, S. & Freckleton, R. P. (2010). Model averaging, missing data and multiple imputation: A case study for behavioural ecology. Behavioral Ecology and Sociobiology 65, 103–116.

Nakagawa, S., Lagisz, M., Jennions, M. D., Koricheva, J., Noble, D. W. A., Parker, T. H., SÁNCHEZ-Tójar, A., Yang, Y. & O’dea, R. E. (2022). Methods for testing publication bias in ecological and evolutionary meta-analyses. Methods in Ecology and Evolution 13, 4–21.

Nakagawa, S., Noble, D. W. A., Senior, A. M. & Lagisz, M. (2017). Meta-evaluation of meta-analysis: Ten appraisal questions for biologists. BMC Biology 15, 18.

Nakagawa, S. & Poulin, R. (2012). Meta-analytic insights into evolutionary ecology: An introduction and synthesis. Evolutionary Ecology 26(5), 1085–1099.

Nakagawa, S. & Santos, E. S. A. (2012). Methodological issues and advances in biological meta-analysis. Evolutionary Ecology 26(5), 1253–1274.

Nakagawa, S., Yang, Y., Macartney, E. L., Spake, R. & Lagisz, M. (2023). Quantitative evidence synthesis: A practical guide on meta-analysis, meta-regression, and publication bias tests for environmental sciences. Environmental Evidence 12, 8.

Navarrete, A. F., Reader, S. M., Street, S. E., Whalen, A. & Laland, K. N. (2016). The coevolution of innovation and technical intelligence in primates. Philosophical Transactions of the Royal Society B: Biological Sciences 371(1690), 20150186.

Noonan, M. P., Sallet, J., Mars, R. B., Neubert, F. X., O’reilly, J. X., Andersson, J. L., Mitchell, A. S., Bell, A. H., Miller, K. L. & Rushworth, M. F. S. (2014). A neural circuit covarying with social hierarchy in macaques. PLoS Biology 12(9), e1001940.

O’donnell, S., Bulova, S., Deleon, S., Barrett, M. & Fiocca, K. (2019). Brain structure differences between solitary and social wasp species are independent of body size allometry. Journal of Comparative Physiology 205(6), 911–916.

O’donnell, S., Bulova, S. J., Deleon, S., Khodak, P., Miller, S. & Sulger, E. (2015). Distributed cognition and social brains: Reductions in mushroom body investment accompanied the origins of sociality in wasps (Hymenoptera: Vespidae). Proceedings of the Royal Society B: Biological Sciences 282(1810), 20150791.

Ott, S. R. & Rogers, S. M. (2010). Gregarious desert locusts have substantially larger brains with altered proportions compared with the solitarious phase. Proceedings of the Royal Society B: Biological Sciences 277(1697), 3087–3096.

Pasquaretta, C., Leve, M., Claidiere, N., van de Waal, E., Whiten, A., MacIntosh, A. J. J., Pele, M., Bergstrom, M. L., Borgeaud, C., Brosnan, S. F., Crofoot, M. C., Fedigan, L. M., Fichtel, C., Hopper, L. M., Mareno, M. C., et al. (2014). Social networks in primates: Smart and tolerant species have more efficient networks. Scientific Reports 4, 7600.

Petit, O., Dufour, V., Herrenschmidt, M., DE Marco, A., Sterck, E. H. M. & Call, J. (2015). Inferences about food location in three Cercopithecine species: An insight into the socioecological cognition of primates. Animal Cognition 18(4), 821–830.

Pick, J. L., Nakagawa, S. & Noble, D. W. A. (2019). Reproducible, flexible and high-throughput data extraction from primary literature: The metaDigitise r package. Methods in Ecology and Evolution 10(3), 426–431.

Preiszner, B., Papp, S., Vincze, E., Bókony, V. & Liker, A. (2015). Does innovation success influence social interactions? An experimental test in house sparrows. Ethology 121(7), 661–673.

R Core Team (2022). R: A language and environment for statistical computing. In R Foundation for Statistical Computing, Vienna, Austria.

Reader, S. M., Hager, Y. & Laland, K. N. (2011). The evolution of primate general and cultural intelligence. Philosophical Transactions of the Royal Society B: Biological Sciences 366(1567), 1017–1027.

Reddy, R. B., Maclean, E. L., Sandel, A. A. & Hare, B. (2015). Social inhibitory control in five lemur species. Primates 56(3), 241–252.

Rehan, S. M., Bulova, S. J. & O’donnell, S. (2015). Cumulative effects of foraging behavior and social dominance on brain development in a facultatively social bee (*Ceratina australensis*). *Brain*, Behavior and Evolution 85(2), 117–124.

Riley, J., Noble, D., Byrne, R. & Whiting, M. (2017). Does social environment influence learning ability in a family-living lizard? Animal Cognition 20, 449–458.

Riley, J. L., Kuchler, A., Damasio, T., Noble, D. W. A., Byrne, R. W. & Whiting, M. J. (2018). Learning ability is unaffected by isolation rearing in a family-living lizard. Behavioral Ecology and Sociobiology 72(2), 1–9.

Rowe, C. & Healy, S. D. (2014). Measuring variation in cognition. Behavioral Ecology 25, 1287–1292.

Sakai, S. T., Arsznov, B. M., Hristova, A. E., Yoon, E. J. & Lundrigan, B. L. (2016). Big cat coalitions: A comparative analysis of regional brain volumes in Felidae. Frontiers in Neuroanatomy 10, 99.

Sakai, S. T., Arsznov, B. M., Lundrigan, B. L. & Holekamp, K. E. (2011). Brain size and social complexity: A computed tomography study in Hyaenidae. Brain Behavior and Evolution 77(2), 91–104.

Sallet, J., Mars, R. B., Noonan, M. P., Andersson, J. L., O’reilly, J. X., Jbabdi, S., Croxson, P. L., Jenkinson, M., Miller, K. L. & Rushworth, M. F. S. (2011). Social network size affects neural circuits in macaques. Science 334, 697–700.

Sandel, A. A., Maclean, E. L. & Hare, B. (2011). Evidence from four lemur species that ringtailed lemur social cognition converges with that of haplorhine primates. Animal Behaviour 81(5), 925–931.

Sandel, A. A., Miller, J. A., Mitani, J. C., Nunn, C. L., Patterson, S. K. & GARAMSZEGI, L. (2016). Assessing sources of error in comparative analyses of primate behavior: Intraspecific variation in group size and the social brain hypothesis. Journal of Human Evolution 94, 126–133.

Schillaci, M. A. (2008). Primate mating systems and the evolution of neocortex size. Journal of Mammalogy 89, 58–63.

Schrijver, N. C. A., Bahr, N. I., Weiss, I. C. & Würbel, H. (2002). Dissociable effects of isolation rearing and environmental enrichment on exploration, spatial learning and HPA activity in adult rats. Pharmacology Biochemistry and Behavior 73, 209–224.

Schubiger, M. N., Fichtel, C. & Burkart, J. M. (2020). Validity of cognitive tests for non-human animals: pitfalls and prospects. Frontiers in Psychology 11, 1835–1835.

Seid, M. A. & Junge, E. (2016). Social isolation and brain development in the ant *Camponotus floridanus*. The Science of Nature 103(5), 42.

Sewall, K. B., Anderson, R. C., Soha, J. A., Peters, S. & Nowicki, S. (2018). Early life conditions that impact song learning in male zebra finches also impact neural and behavioral responses to song in females. Developmental Neurobiology 78(8), 785– 798.

Shaw, R. & Schmelz, M. (2017). Cognitive test batteries in animal cognition research: Evaluating the past, present and future of comparative psychometrics. Animal Cognition 20, 1003–1018.

Shaw, R. C. (2017). Testing cognition in the wild: Factors affecting performance and individual consistency in two measures of avian cognition. Behavioural Processes 134, 31–36.

Shettleworth, S. J. (2001). Animal cognition and animal behaviour. Animal Behaviour 61, 277–286.

Shultz, S. & Dunbar, R. I. M. (2006). Both social and ecological factors predict ungulate brain size. Proceedings of the Royal Society B: Biological Sciences 273(1583), 207– 215.

Shultz, S. & Dunbar, R. I. M. (2007). The evolution of the social brain: anthropoid primates contrast with other vertebrates. Proceedings of the Royal Society B: Biological Sciences 274(1624), 2429–2436.

Shultz, S. & Dunbar, R. I. M. (2010). Social bonds in birds are associated with brain size and contingent on the correlated evolution of life-history and increased parental investment. Biological Journal of the Linnean Society 100, 111–123.

Smith, A. R., Seid, M. A., Jiménez, L. C. & Wcislo, W. T. (2010). Socially induced brain development in a facultatively eusocial sweat bee *Megalopta genalis* (Hailctidae). Proceedings of the Royal Society B: Biological Sciences 277(1691), 2157–2163.

Smith, B. M., Yao, X. Y., Chen, K. S. & Kirby, E. D. (2018). A larger social network enhances novel object location memory and reduces hippocampal microgliosis in aged mice. Frontiers in Aging Neuroscience 10, 142.

Sobrero, R., Fernandez-Aburto, P., LY-Prieto, A., Delgado, S. E., Mpodozis, J. & Ebensperger, L. A. (2016). Effects of habitat and social complexity on brain size, brain asymmetry and dentate gyrus morphology in two Octodontid rodents. Brain Behavior and Evolution 87, 51–64.

Speechley, E.M., Ashton, B.J., Thornton, A., Simmons, L.W. & Ridley, A.R. (2024). Heritability of cognitive performance in wild Western Australian magpies. Royal Society of Open Science 11, 231399.

Stanbrook, E., Jodoin, J., Culbert, B., Shultz, S. & Balshine, S. (2020). Learning performance is influenced by the social environment in cichlid fishes. Canadian Journal of Experimental Psychology 74(3), 215–227.

Sterne, J. A., Gavaghan, D. & Egger, M. (2000). Publication and related bias in meta-analysis: Power of statistical tests and prevalence in the literature. Journal of Clinical Epidemiology 53(11), 1119–1129.

Swanson, E. M., Holekamp, K. E., Lundrigan, B. L., Arsznov, B. M. & Sakai, S. T. (2012). Multiple determinants of whole and regional brain volume among terrestrial carnivorans. PLOS One 7(6), e38447.

Szabo, B., VALENCIA-Aguilar, A., DAMAS-Moreira, I. & Ringler, E. (2022). Wild cognition – linking form and function of cognitive abilities within a natural context. Current Opinion in Behavioral Sciences 44.

Templeton, J. J., Kamil, A. C. & Balda, R. P. (1999). Sociality and social learning in two species of corvids: The pinyon jay (*Gymnorhinus cyanocephalus*) and the Clark’s nutcracker (*Nucifraga columbiana*). Journal of Comparative Psychology 113(4), 450– 455.

Thornton, A. & Lukas, D. (2012). Individual variation in cognitive performance: Developmental and evolutionary perspectives. Philosophical Transactions of the Royal Society B: Biological Sciences 367, 2773–2783.

Todorov, O. S., Weisbecker, V., Gilissen, E., Zilles, K. & Sousa, A. A. D. (2019). Primate hippocampus size and organization are predicted by sociality but not diet. Proceedings of the Royal Society B: Biological Sciences 286, 20191712.

Toyoshima, M., Yamada, K., Sugita, M. & Ichitani, Y. (2018). Social enrichment improves social recognition memory in male rats. Animal Cognition 21(3), 345–351.

Triki, Z., Levorato, E., Mcneely, W., Marshall, J. & Bshary, R. (2019). Population densities predict forebrain size variation in the cleaner fish *Labroides dimidiatus*. Proceedings of the Royal Society B: Biological Sciences 286, 20192108.

Upton, G. (2005). Goodman–Kruskal Measures of Association. In Encyclopedia of Biostatistics (ed. P. Armitage and T. Colton). John Wiley & Sons, Ltd, New Jersey.

VAN Horik, J. & Emery, N. J. (2011). Evolution of cognition. Wiley Interdisciplinary Reviews: Cognitive Science 2(6), 621–633.

Viechtbauer, W. (2010). Conducting meta-analyses in R with the metafor Package. Journal of Statistical Software 36(3), 1–48.

Walker, R., Burger, O., Wagner, J. & VON Rueden, C. R. (2006). Evolution of brain size and juvenile periods in primates. Journal of Human Evolution 51(5), 480–489.

Wang, L., Cao, M., Pu, T., Huang, H., Marshall, C. & Xiao, M. (2018). Enriched physical environment attenuates spatial and social memory impairments of aged socially isolated mice. International Journal of Neuropsychopharmacology 21(12), 1114– 1127.

Wascher, C. A. F. (2015). Individual performance in socio-cognitive tasks predicts social behaviour in carrion crows. Behaviour 152(5), 615–634.

Weisbecker, V., Blomberg, S., Goldizen, A. W., Brown, M. & Fisher, D. (2015). The evolution of relative brain size in Marsupials Is energetically constrained but not driven by behavioral complexity. Brain Behavior and Evolution 85(2), 125–135.

Weiss, I. C., Pryce, C. R., Jongen-Rêlo, A. L., Nanz-Bahr, N. I. & Feldon, J. (2004). Effect of social isolation on stress-related behavioural and neuroendocrine state in the rat. Behavioural Brain Research 152(2), 279–295.

Wheeler, D. L., Barrett, T., Benson, D. A., Bryant, S. H., Canese, K., Chetvernin, V., Church, D. M., Dicuccio, M., Edgar, R., Federhen, S., Geer, L. Y., Helmberg, W., Kapustin, Y., Kenton, D. L., Khovayko, O., et al. (2006). Database resources of the National Center for Biotechnology Information. Nucleic Acids Research 34, D173–D180.

Williams, B. M., Luo, Y., Ward, C., Redd, K., Gibson, R., Kuczaj, S. A. & Mccoy, J. G. (2001). Environmental enrichment: Effects on spatial memory and hippocampal CREB immunoreactivity. Physiology and Behavior 73(4), 649–658.

